# Reconstructing colonization dynamics to establish how human activities transformed island biodiversity

**DOI:** 10.1101/2023.02.09.526923

**Authors:** Sean Tomlinson, Mark Lomolino, Sean Haythorne, Atholl Anderson, Jeremy J. Austin, Stuart Brown, George Perry, Janet Wilmshurst, Jamie R. Wood, Damien A. Fordham

## Abstract

**Aim:** The drivers and dynamics of initial human migrations across individual islands and archipelagos are poorly understood, affecting assessments of human-modification of island biodiversity. Here, we describe and test a process-explicit approach for reconstructing human arrival and expansion on islands, which combines archaeological and climate records with high-resolution spatial population models. Using Polynesian colonisation of New Zealand as an example, we show that our new method can generate information crucial for assessing how humans affected biodiversity on islands.

**Innovation:** The transition of islands from prehuman to human dominated ecosystems has typically been assessed by comparing biodiversity before and after time of first arrival, without considering the potential importance of the spatiotemporal dynamics of the human expansion event. Our new approach, which uses pattern-oriented modelling methods to combine inferences of human colonisation dynamics from dated archaeological material with spatially explicit population models, produces validated reconstructions of the pattern and pace of human migration across islands at high spatiotemporal resolutions. From these reconstructions, demographic and environmental drivers of human colonization can be identified, and the role that people had on biodiversity established. Using this technique, we show that closely reconciling inferences of Polynesian colonisation of New Zealand requires there to have been a single founding population of approximately 500 people, arriving between 1233 and 1257 AD, settling multiple areas, and expanding quickly over both North and South islands. The resultant maps of Māori colonisation dynamics provide new opportunities to better determine how human activities transformed biodiversity of New Zealand in space and time.

**Main conclusions:** Process-explicit models can reconstruct human migration across large islands, producing validated, high resolution spatiotemporal projections of human occupancy and abundance that account for dispersal and population dynamics. This modelling framework should prove effective across any islands and archipelagos where climate and archaeological records are available.

## Introduction

The emergence of hominids and their sequential dispersal away from an African evolutionary cradle has always been an intriguing topic for biogeographers, ecologists and conservation biologists (Diamond, 1997; Finlayson, 2005; Wallace, 1876). However, key questions remain concerning the timing, rate and mechanisms influencing the rapid expansions of our species (Mellars, 2006; Nielsen et al., 2017) and the broader ecological consequences of human colonization on biodiversity (Burney & Flannery, 2005; Ellis, 2021). This is particularly true for the colonisation of remote oceanic islands (Channell & Lomolino, 2000; Russell & Kueffer, 2019; Wood, 2008), which are among the last areas on Earth to have been settled and transformed by people (Nogué et al., 2021).

While several potential pathways for the global expansion of modern humans have been proposed (Beyer, Krapp, Eriksson, & Manica, 2021; Eriksson et al., 2012; Timmermann & Friedrich, 2016), simulations of these colonisation dynamics have, to date, been done at relatively coarse spatiotemporal scales, often underpinned by an assumed positive correlation between net primary productivity and population growth in pre-agricultural societies (Zhu, Galbraith, Reyes-García, & Ciais, 2021). Consequently, knowledge of drivers of human migration and their fine-scale dynamics are unclear, particularly for those that operated at spatial and temporal scales relevant to individual islands and archipelagos (Douglass et al., 2019; Wilmshurst, Anderson, Higham, & Worthy, 2008).

Human arrival dates on many islands and archipelagos have been established archeologically with reasonable certainty (Wilmshurst, Hunt, Lipo, & Anderson, 2011). While these dates have often been used to speculate on the role and impact of human activities on island biodiversity (Duncan & Blackburn, 2004; Nogué et al., 2021; Wood et al., 2017), this has typically been done without considering the additional and important roles that founding population size and location, and rate and pace of expansion could have had on the spatiotemporal pattern of biodiversity. This oversight has not been intentional, but rather has occurred because of an absence of high-resolution reconstructions of human migrations across islands, which is difficult to establish, and remains heavily contested for most islands (Hansford, Lister, Weston, & Turvey, 2021; Rieth, Hunt, Lipo, & Wilmshurst, 2011; Walter, Buckley, Jacomb, & Matisoo-Smith, 2017).

Improving knowledge of the processes responsible for the transformation of native insular biotas following human arrival and expansion requires new methods that can reconstruct human colonization dynamics at spatiotemporal resolutions required for biodiversity assessments. These include assessments of the causal role of people on extinctions of island endemics, and resultant changes in the ecological function of islands across the Pacific Ocean (Boyer & Jetz, 2014), Indian Ocean (Anderson et al., 2018; Hixon et al., 2021; Wood et al., 2017), the Mediterranean (Wood et al., 2017), and the Caribbean (Cooke, Dávalos, Mychajliw, Turvey, & Upham, 2017; Locatelli, Due, van den Bergh, & van den Hoek Ostende, 2012). New methods in macroecology that synthesize disparate evidence from archaeological records have the potential to reconstruct human events at spatiotemporal resolutions requisite for establishing human-mediated biodiversity change on islands (Fordham et al., 2020). However, their application to island systems has yet to be tested.

Part of the challenge with spatiotemporally reconstructing the dynamics of initial human migration across individual islands and archipelagos is that most remote islands were settled rapidly and relatively recently, when climates were similar to current conditions (Nogué et al., 2021). Consequently, these events cannot be reconstructed adequately in space and time using existing correlative techniques (Giampoudakis et al., 2017), or climate proxies (Beyer et al., 2021). A potential solution could be to integrate archaeological information with spatially explicit population models (SEPM) that can reconstruct fine-scale dispersal and population dynamics using process-explicit approaches and pattern-oriented methods (Fordham, Haythorne, Brown, Buettel, & Brook, 2021). Process-explicit approaches simulate the dynamics of a biological system as explicit functions of the events that drive changes in that system (Pilowsky, Colwell, Rahbek, & Fordham, 2022). When coupled with pattern-oriented modelling (POM) approaches (Grimm & Railsback, 2012) — an emerging and powerful tool in macroecology and biogeography (Honkaniemi, Rammer, & Seidl, 2021) — process-explicit models can establish chains of causality responsible for colonisation and extinction dynamics (Fordham et al., 2022), and resultant biodiversity change (Rangel et al., 2018). Critically, the approach has substantial potential for reconstructing rapid human expansion at relatively fine spatial scales, including those across oceanic islands and archipelagos during periods of climatic stasis (Fordham et al., 2021).

The Māori expansion across New Zealand provides an intriguing and insightful model system to demonstrate how the colonisation and subsequent expansion dynamics of humans across islands can be reconstructed using an approach that combines SEPMs (Wiegand, Moloney, Naves, & Knauer, 1999) with POM methods (Grimm & Railsback, 2012). This is because there is a wealth of precisely dated archaeological evidence of Māori activity (S. J. Holdaway et al., 2019), existing models of human population growth (Brown & Crema, 2019; R. N. Holdaway et al., 2014; R. N. Holdaway & Jacomb, 2000), and 18^th^ century estimates of population size (Pool, 1991). Just as importantly, there is an immediate need for a more detailed understanding of the pattern and pace of Māori migration across New Zealand to better understand the role past human activities had on the dynamics and extinctions of New Zealand’s native biotas. This is because current assessments of biodiversity change following the peopling of New Zealand have rarely considered the consequences of founding location, or rate and pattern of human expansion across the archipelago (R. N. Holdaway & Jacomb, 2000; G. L. W. Perry, Wilmshurst, McGlone, & Napier, 2012).

The East Polynesian expansion in the Pacific Ocean was the final phase of global human settlement (Wilmshurst et al., 2008). It included the colonisation of the New Zealand archipelago by Polynesians known subsequently as Māori. Archaeological evidence suggests an expansion that was so rapid as to appear highly synchronous across the entire archipelago (Anderson, 1991; Walter et al., 2017). Consequently, there remains little consensus on the location of first arrival, migration routes and whether colonisation resulted from a series of small founding populations or a single, concerted migration (Anderson, 2017; Walter et al., 2017; Wilmshurst et al., 2008): information urgently needed to better understand the human dimension of biodiversity change in New Zealand.

These colonization dynamics cannot be resolved using existing human-migration models, partly because they rely on climatic change (and derived changes in net primary productivity) as the principal drivers of colonisation and expansion (Eriksson et al., 2012; Zhu et al., 2021). However, climatic conditions in New Zealand during the period of colonisation (1200 – 1300 AD) were relatively stable (Wanner et al., 2008), providing no insights into the establishment and spread of people across New Zealand, nor their subsequent spatiotemporal impacts on native biotas. Moreover, Polynesian colonists brought with them horticulture (but see Anderson and Petchey (2020)) and, thus, were not entirely dependent on hunting and gathering (Anderson, 2016; Brown & Crema, 2019; Furey, 2006), for which net primary productivity is a proxy (Zhu et al., 2021). It is likely, however, that these limitations can be overcome using process-explicit models, archaeological records and climate and environmental data (Fordham et al., 2020).

The process-explicit, pattern-oriented modelling framework that we develop and test here, simulating the colonization of New Zealand, has great potential for understanding how Māori transformed island biodiversity. More generally, it can be used to reconstruct the initial waves of human colonisation across other remote large islands and archipelagos, potentially providing novel insights into fine-resolution drivers of biodiversity change following human arrival.

## Methods

Our new spatially explicit population modelling (SEPM) approach for reconstructing human colonisation dynamics on islands at high spatiotemporal resolutions integrates archaeological data with population growth and dispersal models to produce dynamic simulations of changing populations, distributions and migration routes of people at fine spatiotemporal resolutions (Figure 1). Archaeological records matched with climate and environmental data are used to reconstruct habitat suitability for humans on islands, and relative density patterns at spatial resolutions required to capture local orographic influences (Supplementary Figure 1). This information is integrated into SEPMs that simulate population growth and dispersal dynamics. Uncertainty is captured directly in simulations by varying model parameters (demographic, dispersal, suitability, and density parameters), producing thousands of conceivable models of human arrival and establishment (Figure 1). Pattern-oriented modelling (POM) methods are used to optimise parameter values using inferences of demographic change from archaeological and historical records. Models that validate well are used to reconstruct human colonisation and establishment, and to identify causative processes responsible for spatiotemporal patterns, generating information needed to determine past influences of people on biodiversity (Fordham et al., 2022). Resultant conclusions can be tested using counterfactual scenarios that modify the effects of these parameters (G. L. Perry, Wainwright, Etherington, & Wilmshurst, 2016).

**Figure 1:**
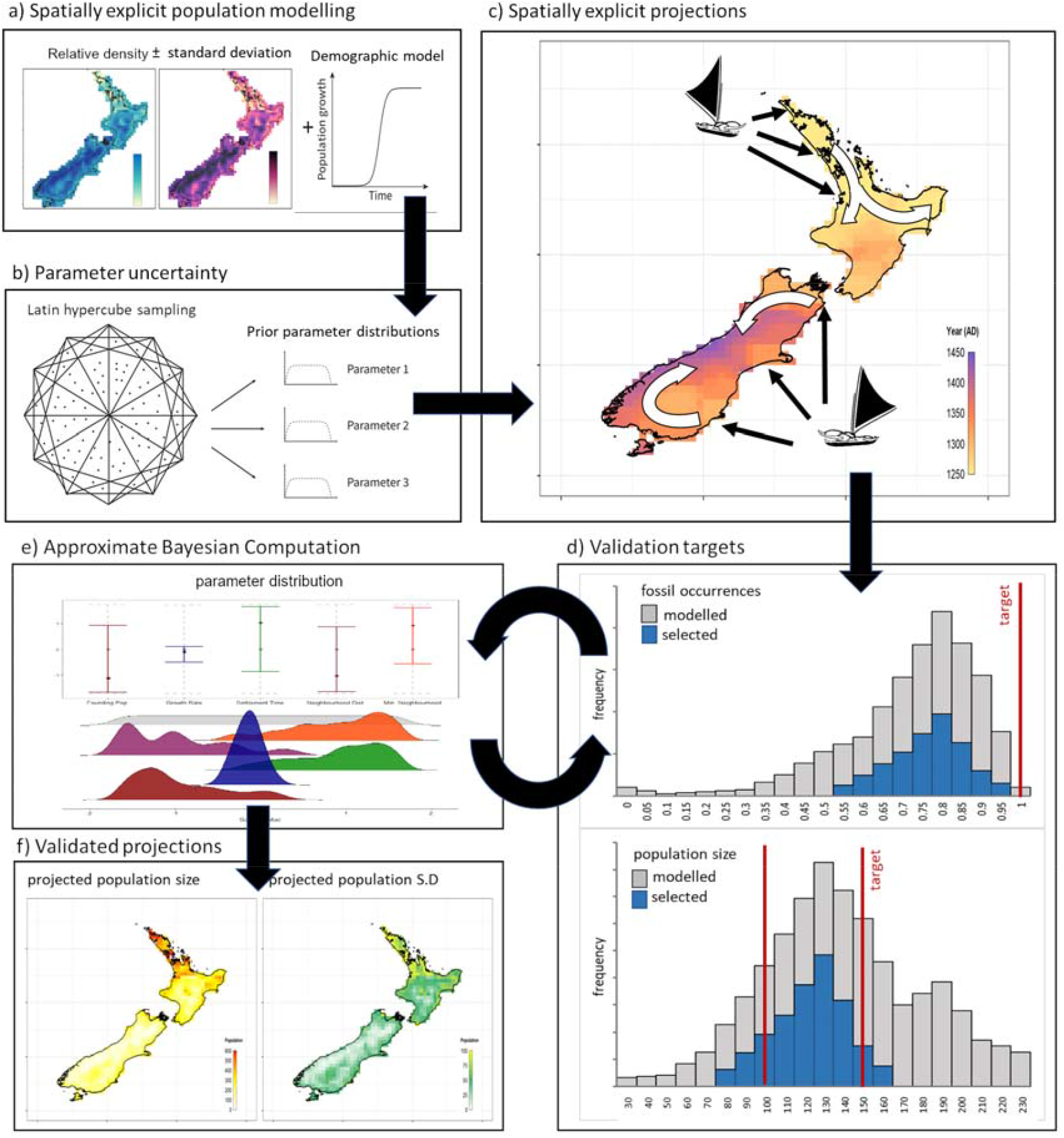
Reconstructing Māori colonisation dynamics using spatially explicit population modelling (SEPM) with pattern-oriented modelling (POM). **(a)** Spatiotemporal estimates of Māori relative density were combined with demographic models to simulate colonisation, population growth and geographic expansion. **(b)** To account for parameter uncertainty, thousands of potential models were generated using Latin hypercube sampling, and **(c)** each model was simulated, providing a plausible spatiotemporal projection of arrival time, range expansion and population abundance. **(d)** Model projections were validated against inferences from archaeological archives, and **(e)** the ‘best’ projections were selected using Approximate Bayesian Computation. The frequency distribution of parameters in these best models were compared to their frequency distribution for all models, and if they differed the processes was repeated. **(f)** Once the parameters converged, the best models were used to project population abundance in space and time

Below we describe the application of this approach to the colonization of New Zealand by Māori. Models are coded in Program R version 4.0.4 (R Core Team, 2021) and are described in detail in the Supporting Methods, and example simulations which are available here: https://figshare.com/s/02c292e2386633546e2e.

### Modelling Māori relative population density

Spatial models of relative density of Māori populations prior to European first contact (conventionally 1769 C.E.) can be constructed using the density of archaeological finds as a proxy for human density (Goldberg, Mychajliw, & Hadly, 2016). Specifically, we trained boosted regression tree models (BRT; Elith, Leathwick, and Hastie (2008)) using radiocarbon (^14^C) dated Māori archaeological sites from the New Zealand Radiocarbon database (Figure 2), which we intersected with paleoclimate data generated using PaleoView v1.5.1 (Fordham et al., 2017), and geomorphometric data. We used a decomposed hurdle approach for BRT models (Ridout, Demétrio, & Hinde, 1998). This allowed the occurrence and abundance of ^14^C data to be trained on different environmental factors (Potts and Elith (2006); Ridout et al. (1998); Supplementary Figure 2), and addressed zero inflation in the archaeological database (Mellin, Russell, Connell, Brook, & Fordham, 2012). Zero inflation in the database resulted from both a spatial absence of archaeological finds, and from a lack of ^14^ C dated artefacts at some archaeological sites. The BRT model was used to project mean relative population abundance and its standard deviation across New Zealand at a grid cell resolution of 0.25° x 0.25° (Figure 2). These projections were used as a spatial template for the SEPM. The BRT model and its validation are described in detail in the Supporting Methods.

**Figure 2:**
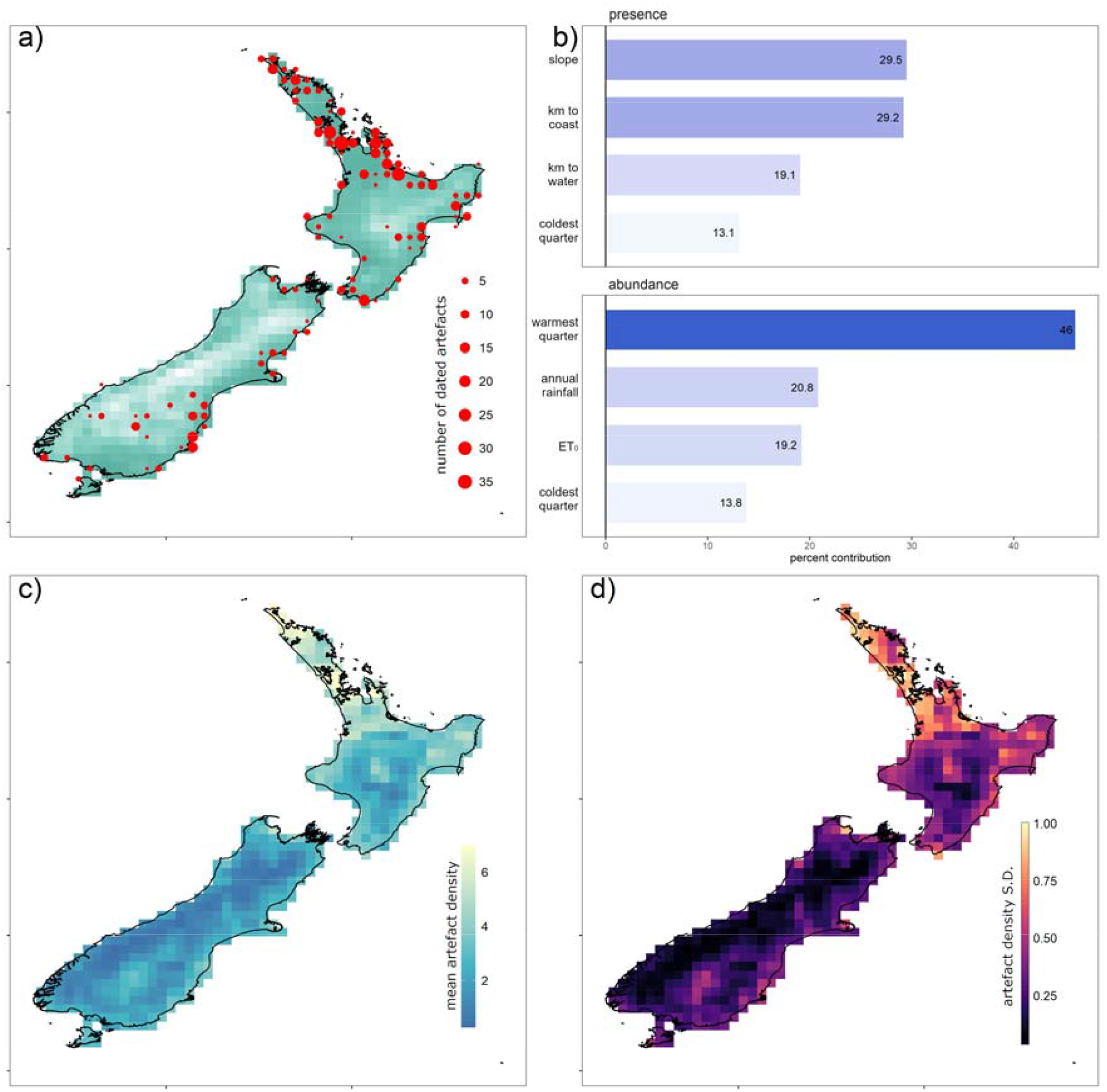
Reconstructing relative abundance of people using archaeological data. **(a)** ^14^C dated archaeological artefacts across New Zealand during the colonisation period (1000 to 1650 C.E.) mapped at a 0.25° resolution. Lighter cells represent higher elevations. **(b)** Effect sizes for variables contributing to the probability of presence and the relative abundance of human artefacts (proxies for presence and abundance of people) across the period of colonisation in New Zealand (estimated using a boosted regression tree). **(c)** Map of mean relative density of human artefacts and **(d)** its standard deviation. Together **(c)** and **(d)** form the spatial template for the spatially explicit population models (SEPMs). Variables in **(b)** are the area of each grid cell steeper than 20° (slope), the distance to the coast (km to coast), the distance to navigable water (km to water), the average temperatures in the coldest quarter (coldest quarter) and the warmest quarter (warmest quarter) of the year, annual rainfall, and annual evapotranspiration (ET_0_)

### Spatially Explicit Population Model (SEPM)

To reconstruct the colonisation dynamics of Māori from 1230 – 1850 AD, spatial projections of potential relative population abundance and its standard deviation (described above) can be coupled with a population growth model and a dispersal simulator (Figure 3). This is done using a lattice-based SEPM framework (Fordham et al., 2021) that models range expansion annually as a function of population size (Supplementary Figure 3) and habitable neighbourhoods (Figure 3). To do this for Māori, we used an existing exponential population growth model (R. N. Holdaway et al., 2014; R. N. Holdaway & Jacomb, 2000), colonising neighbourhoods sequentially. Neighbourhoods with the highest potential relative abundances (those in the most suitable areas) were colonised first. To do this, grid cells were grouped into spatial neighbourhoods using foraging radii. This allowed dispersal of the Māori population across New Zealand to be simulated as the total population grew (Figure 3).

**Figure 3:**
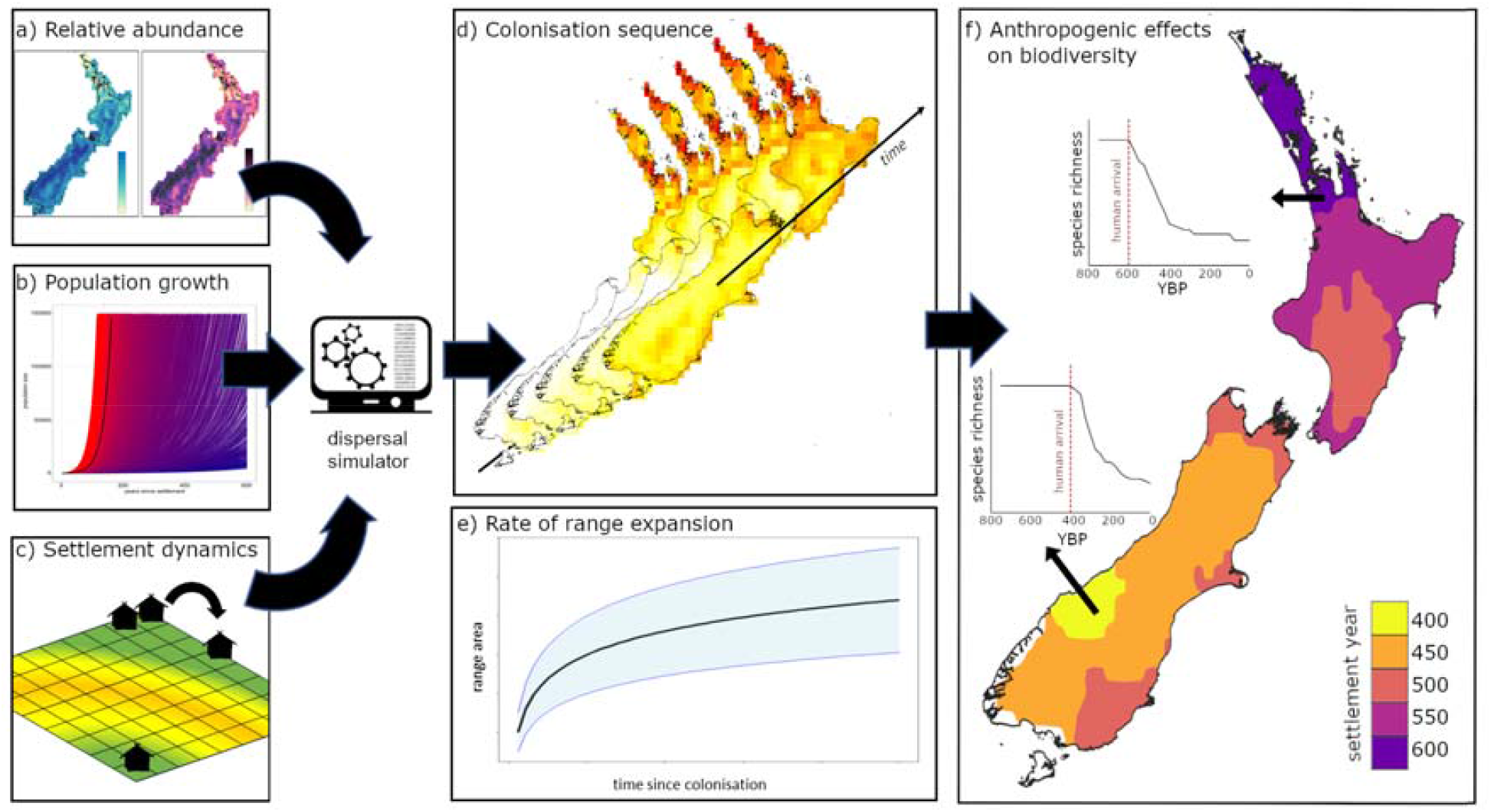
Simulating range expansion using spatially explicit population model (SEPMs). Māori migration is simulated using **(a)** statistical estimates of relative abundance (see Figure 2), **(b)** demographic growth models and **(c)** the minimum number of people required to found a new community. This results in **(d)** spatiotemporal estimates of timing of colonisation and **(e)** temporal estimates of rate of range expansion. These results can be mapped as **(f)** estimates of the timing of human arrival, establishment and growth: information required to establish the role of human activities on island biodiversity change, including changes in species richness (as indicated here), ranges of species and ecosystem structure.

To account for parameter uncertainty, we generated 25,000 potential simulations. We did this by varying five parameters in the SEPM across large but plausible ranges (Table 1) using a robust coverage of multi-dimensional parameter space (Fordham, Haythorne, & Brook, 2016). Variable parameters were time of arrival, founding population, population growth rate, neighbourhood size, and foraging distance. The SEPM was built using the ‘poems’ version 1.0.1 Program R package (Haythorne, Fordham, Brown, Buettel, & Brook, 2021). A detailed description of the mechanics of the model is provided in the Supporting Methods.

**Table 1:** Parameter values used in the process-explicit model (POM) of Māori colonisation and expansion. Fixed values were consistent across all simulations, while variable parameters were allowed to vary randomly across the entire parameter space (Type). Fixed and prior values for parameters are provided (Values). Posterior values for variable parameters according to POM validation are shown (Credible Interval). Superscript letters indicate published sources for credible intervals; a = (R. N. Holdaway & Jacomb, 2000); b = (G. L. Perry et al., 2014); c = (Wilmshurst et al., 2008).

| *poems* parameter | Description | Type | Values | Credible interval |
| --- | --- | --- | --- | --- |
| ***Spatiotemporal template*** |  |  |  |  |
| simulation_start_year | The year at which all simulations were initiated | fixed | 850 AD | NA |
| time steps | Simulation years | fixed | 1101 | NA |
| Lattice | Number of 0.25° grid-cells | fixed | 431 | NA |
| ***Demographic parameters*** |  |  |  |  |
| human_founding_population | The number of Polynesian colonists that first arrived in New Zealand | variable | 200 - 600 | 435-582<sup>a</sup> |
| human_growth_rate | The rate at which the Māori population increased following arrival in New Zealand | variable | 1.005 – 1.050 | 1.010-1.011<sup>a</sup> |
| ***Settlement and expansion parameters*** |  |  |  |  |
| human_colonisation_time | The year (AD) in which Polynesians colonised New Zealand | variable | 1230 - 1314 | 1233-1257<sup>b, c</sup> |
| human_foraging_distance | The radius (km) of the foraging range of each Māori settlement | variable | 30 – 70 | 63-68 |
| human_min_neighbourhood | The minimum number of people required to seed a new Māori settlement | variable | 20 – 80 | 23-36 |

### Pattern oriented Model Validation

POM methods can be used to optimise parameters in SEPMs (Haythorne et al., 2021). This is done by comparing model simulations with independent validation targets and selecting models that have the mechanisms to most closely replicate these targets (Grimm & Railsback, 2012), often using Approximate Bayesian Computation (ABC; Csilléry, Blum, Gaggiotti, and François (2010)).

Model simulations of Māori arrival and expansion in New Zealand were assessed against two targets: (i) Spatiotemporal occurrence, measured as modelled presence in grid cells at a time and place where ^14^C-dated archaeological evidence indicated that that the grid cells should have been occupied; and (ii) Population size, measured as a population of between 100,000 and 150,000 people across the archipelago in 1769 C.E. based on the earliest estimate made by the British explorer Captain James Cook, including its likely uncertainty (Pool, 1991). The best 1 % of simulations were selected using the rejection algorithm in the ‘*abc*’ package (Csilléry, François, & Blum, 2012). The parameter ranges identified by ABC as most accurately matching the validation targets were used to build additional simulation models (n = 25,000), using the posteriors of previous model runs as informed priors (Pilowsky et al., 2022). This POM process was stopped when Bayes factors indicated that the selected posteriors no longer differed from the informed priors (Gelman, Hwang, & Vehtari, 2014). Posterior predictive checks were used to determine whether the posterior distributions generated strong resemblance between the simulation results and observed data (Gelman et al., 2014). See the Supporting Methods for further details.

### Model output and Sensitivity analysis

To reconstruct human colonization patterns, we calculated credible intervals for model parameters from the ‘best’ 1 % of optimised simulations and then generated multi-model averaged projections of time and location of first arrival of Māori in New Zealand, founding population size, and population growth and migration through space and time. Projections were weighted by ABC model weights (Fordham et al., 2022).

We determined the sensitivity of the results to two common model-based structural assumptions (Saltelli, Tarantola, & Campolongo, 2000): the form of the population growth model; and the number of founding events. To do this, 25,000 simulations were generated using a robust coverage of the posterior parameter space identified by the POM, altering human growth so that it followed a logistic, rather than exponential, function (Brown & Crema, 2019); and by making founding events > 1 (i.e., multiple rather than a single fleet). Where multiple founding events were simulated, founding populations were spread over multiple time steps. Model outputs were compared to simulations without these structural changes. See Supporting information for further details.

## Results

### Māori relative population density

The likelihood of the occurrence of Māori was higher in areas with fewer steep slopes (i.e. > 20°) and those closer to navigable waters (Figure 2; Supplementary Figure 2). Higher relative densities of Māori were projected to occur in areas where average temperatures during the warmest three months of the year exceeded 18 °C, temperatures in the coldest three months exceeded 10 °C, where evapotranspiration (and thus horticultural productivity) was high, and where rainfall was limited, preventing water logging of crops (Figure 2; Supplementary Figure 2). The reconstructed pattern of Māori density strongly replicated the distribution of archaeological sites, with 90% of archaeological sites having > 0.75 likelihood of Māori occupancy.

### Colonisation dynamics

Reconstructing validation targets for human colonisation of New Zealand required a constrained set of ecological parameters: a founding population size of 517 (min: 435 to max: 582), a colonisation year of 1244 C.E. (1233-1257), a minimum community size of 28 individuals (23-36), a neighbourhood radius of 66 km (63-68), and a population growth rate of 1.010 per annum (1.010-1.011) (Table 1, Figure 4). While the first iteration of POM (with broad uninform priors) resulted in selected models that replicated colonisation patterns reasonably well (Figure 4), the second and third iterations of POM did better, placing colonists at nearly all known settlements prior to the earliest radiocarbon dated evidence of their presence there (Figure 4). The best models of the third iteration yielded estimates of population size in 1769 [119,900 (88,750-159,197)] that most closely matched the target (Figure 4).

**Figure 4:**
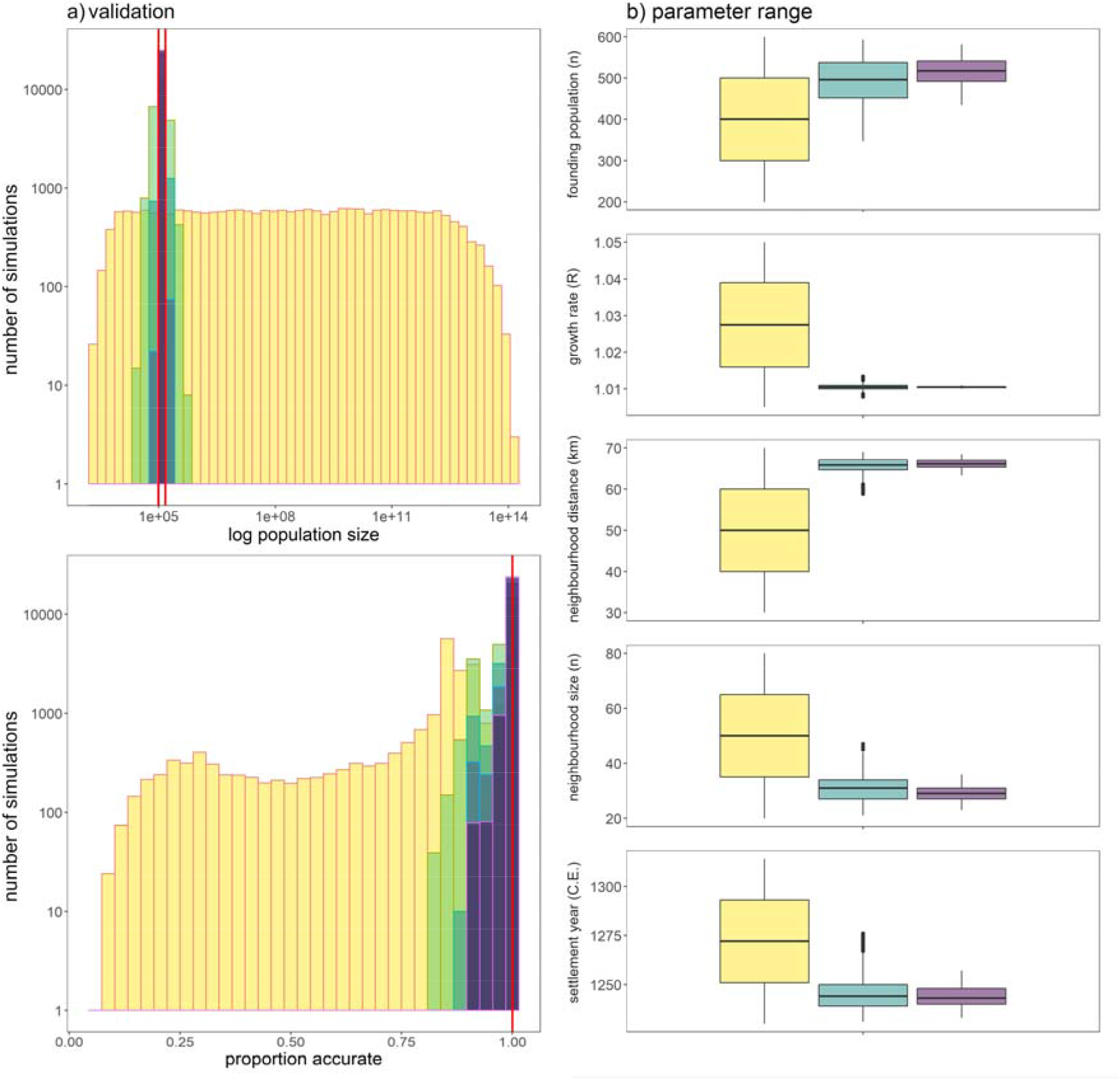
Estimates of settlement and colonisation of New Zealand by Māori using pattern-oriented modelling (POM). **(a)** Histograms show differences between simulated frequencies and observed targets for three iterations of the model, refined using Approximate Bayesian Computation (ABC). Top histogram shows results for population size at time of first European contact (plotted on the log scale). Bottom histogram shows the proportion of archaeological records accurately modelled in space and time. Red solid lines indicate validation targets. **(b)** Box plots show ranges for model parameters resulting from reiterative ABC resampling. In **(a)** and **(b)**, yellow represents the first iteration, green indicates the second iteration, and purple indicates the third iteration.

The best 1 % of SEPMs (from the third POM iteration) consistently simulated the North Island being colonised prior to the South Island (Figure 5). They simulated the colonisation of New Zealand as occurring rapidly, with the entirety of habitable regions colonised by approximately 1400 C.E.; i.e., within 200 years of arrival (Figure 5; Supplementary animation 1). On the South Island, present day Otago, Canterbury, Marlborough and Nelson were projected to have been settled as early as the mid-1200s C.E. in some selected simulations (Figure 5; Supplementary Figure 1). While small differences in settlement and dispersal parameters between selected simulations caused some variation in reconstructions of occupancy and abundance, there was substantial spatiotemporal agreement between the best 1 % of simulations for the pattern of Māori establishment of New Zealand (Supplementary animation 2). Based on the multi-model average of selected models, approximately 63% of the Māori population lived in areas of present-day Northland, Auckland, Waikato, Taranaki and Bay of Plenty during the colonising period, a finding consistent with earlier suggestions that these regions harboured the largest Māori populations (Brown & Crema, 2019; M. McGlone, Anderson, & Holdaway, 1994). Areas of fastest population growth occurred across the North Island, especially in present day Wairarapa, Manawatu and Wellington, and in Nelson and Canterbury on the South Island (Figure 5; Supplementary Figure 1).

**Figure 5:**
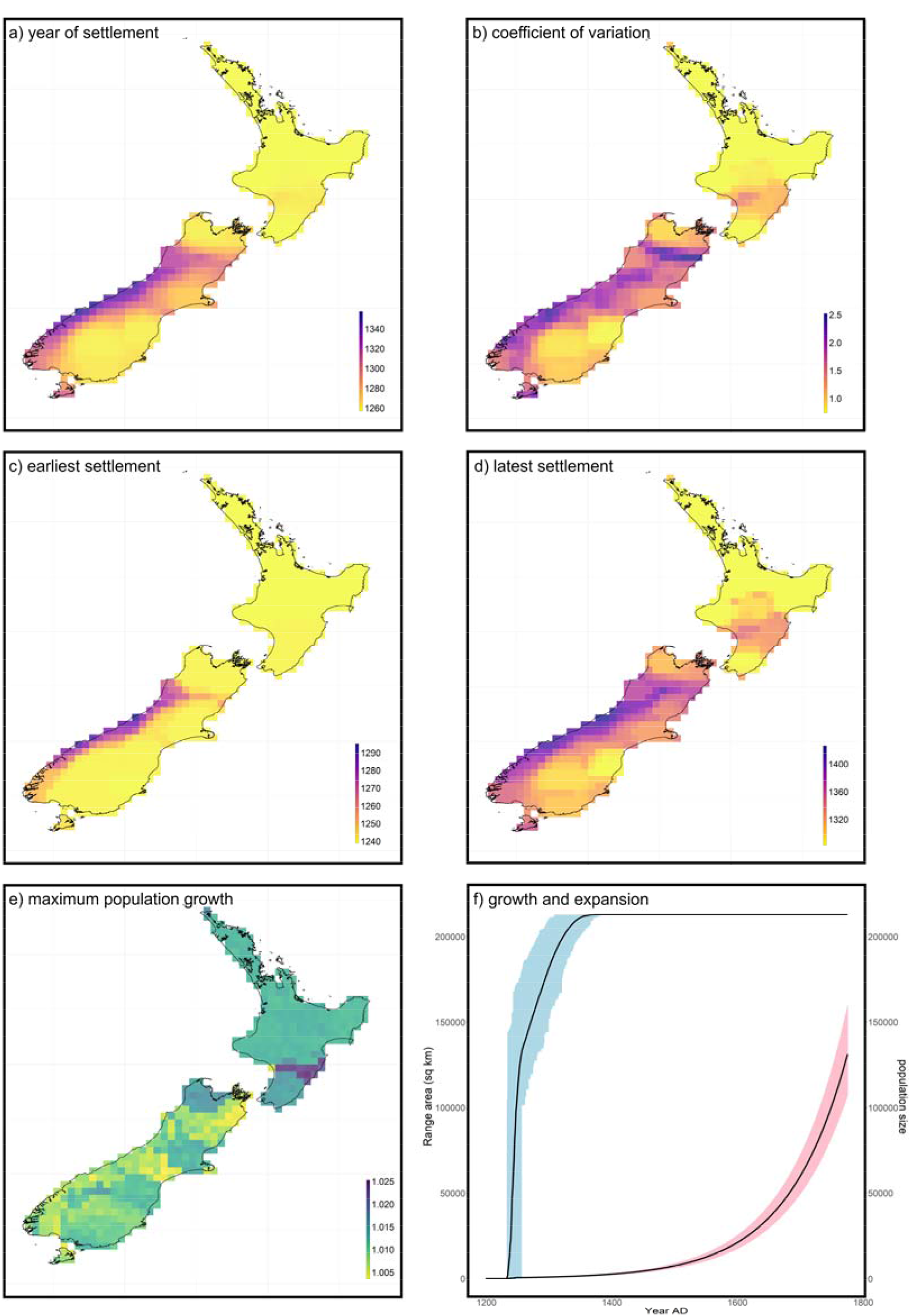
Island colonisation dynamics. Spatial estimates of **(a)** mean colonisation year and **(b)** its coefficient of variation, **(c)** earliest and **(d)** latest estimate of colonisation, and **(e)** maximum population growth rate. Estimates are multi-model ensemble average based on the 250 spatially explicit population models that best reconciled validations targets. **(f)** Estimated rate of range expansion (blue) and population growth (pink). The ensemble mean is shown in black.

Our projections of Māori arrival and expansion in New Zealand were not sensitive to the population growth function (i.e., logistic versus exponential; Supplementary Figure 6). However, the number of founding events substantially altered the pattern and timing of colonisation, and these differences were magnified with increasing numbers of founding events (Supplementary Figure 7). Total population size in the year 1769 C.E. was sensitive to both the type of population growth function and number of founding events (Supplementary Figures 6 and 7). The implications of these observations are discussed below.

## Discussion

Given the integral role that human population growth and expansion has had on biodiversity declines during the Holocene (Burney & Flannery, 2005; Channell & Lomolino, 2000), understanding how humans colonised different islands and archipelagos in response to their unique environments is key to understanding the ecological consequences of these events, including globally-significant declines in biodiversity (Nogué et al., 2021). However, absence of high-resolution reconstructions of patterns and paces of human migrations across islands continues to hinder the extent to which islands can be used as replicated model systems to establish processes of human transformation of biodiversity. We show that process-explicit models that are informed by the archaeological record and spatiotemporal reconstructions of past climates and environments can provide new and important insights into the patterns and drivers of colonisation and establishment of people on islands, generating spatiotemporal reconstructions of human abundance at resolutions needed for biodiversity assessments.

Our SEPM projections of the arrival and expansion of Māori in New Zealand closely reconciled inferences of demographic change from the archaeological record, and more recent historical observations, revealing the importance of topography, proximity to navigable water bodies, and the geography of climatic conditions and habitats on colonisation dynamics. These validated simulations provide new opportunities to explore more extensively the potential ecological impacts of human colonisation on New Zealand’s native biota and ecosystems in space and time (Greig & Rawlence, 2021; M. S. McGlone & Wilmshurst, 1999; G. L. Perry, Wheeler, Wood, & Wilmshurst, 2014), including the roles people have had on species distributions and changes in species richness and ecological function (Figure 3). More generally, the framework developed for reconstructing the colonization of New Zealand by Māori, is directly transferrable to other islands and archipelagos, where climate and archaeological records are available.

### Spatially Explicit Insights on Māori Colonisation

While Polynesian expansion across the Pacific is hypothesised to have resulted from carefully planned, specific colonial intentions (Diamond, 1985), others have argued otherwise (Anderson, Binney, & Harris, 2015; Walter et al., 2017). Our modelling supports the assertion of Diamond (1985), showing that Polynesian colonisation of New Zealand was highly synchronous, with early settlements arising nearly simultaneously in multiple locations, probably connected by coastal navigation routes. Parameter values in our models, chosen through pattern-oriented methods, are highly congruent with established estimates, including timing of arrival in New Zealand (G. L. Perry et al., 2014; Wilmshurst et al., 2008), number of colonisers (Anderson, 2017; Walter et al., 2017; Whyte, Marshall, & Chambers, 2005), and spatial variation in population growth rates (Brown & Crema, 2019). The areas projected by our models as the most likely sites of Māori first settlement also encompass sites with the oldest archaeological evidence of Māori presence, such as Wairau Bar, Houhora and Tairua (R. N. Holdaway et al., 2014; Kinaston et al., 2013; Wilmshurst, Higham, Allen, Johns, & Phillips, 2004)

Sensitivity analysis indicated that increasing the number of independent founding events above one substantially altered the projected colonisation dynamics, resulting in a poorer match between model simulations and inferences of demographic change from the archaeological record. This suggests that New Zealand was likely to have been founded by a single colonisation event. However, this result must be viewed cautiously since the parameters of models with founding events greater than one were not optimised using POM approaches (Pilowsky et al., 2022). Nevertheless, a very high (and perhaps unrealistic) population growth rate would be needed to reproduce the archaeological record under a scenario of multiple founding events.

An element of the Māori colonisation of New Zealand that we could not replicate was the putative abandonment of the South Island following the extinction of the moa, which has been inferred from the fossil record (Rawlence et al., 2015; Walter et al., 2017). Some authors have suggested that the South Island was never densely populated by Māori (Diamond, 1997), as indicated by our SEPM, and that sparse populations persisted following the depletion of wild food resources such as moa (Brown & Crema, 2019; Hamel, 1982). However, this runs contrary to the prevailing view that the South Island initially harboured large Māori populations who then shifted to the North Island when wild food sources were depleted (Rawlence et al., 2015; Walter et al., 2017).

### Ecological Implications of Rapid Colonisation

The arrival and spread of humans across the world’s islands had substantial ecological consequences (Russell & Kueffer, 2019), and the Polynesian colonisation of New Zealand was no different. The colonisation of New Zealand resulted in widespread deforestation (G. L. W. Perry et al., 2012), wholesale extinctions of the terrestrial megafauna (R. N. Holdaway & Jacomb, 2000; G. L. Perry et al., 2014), and serious declines in marine mammal populations (Smith, 2013). However, until now, the timing, rate and magnitude of these anthropogenic impacts have been difficult to resolve because of the absence of a detailed spatiotemporal understanding of how Māori expanded across the archipelago.

Our new macroecological modelling approach for reconstructing the peopling of islands shows strong spatiotemporal variation in colonisation patterns of New Zealand and subsequent densities of people. We project that colonisation happened more rapidly on the North Island, spreading from the northwest of the island to the southeast. On the South Island the colonisation and spread of people is likely to have happened more slowly, spreading from the east of the island to the west. Given that human density and environmental change are strongly correlated at local-to-regional scales (Ellis, 2021), this fresh perspective on Māori colonisation dynamics is likely to provide important new insights into the ecological impacts of this rapid migration of humans across New Zealand.

Although our modelling shows that Māori are likely to have had little ecological impact on the forests west of the Southern Alps, the pervasive impacts of altered fire regimes (G. L. W. Perry et al., 2012) and introduced commensals such as the kiore (*Rattus exulans*) or the kurī (*Canis familiaris*) were significant (Greig & Rawlence, 2021; Wilmshurst et al., 2008). Accordingly, future modelling exercises that investigate biodiversity change following human-colonisation of New Zealand will ideally need to include the likely impacts of commensals and their cascading effects on native, insular biota.

### Broader application

While New Zealand presents a tractable example of human colonisation and expansion, resulting in a globally-significant decline in biodiversity (Duncan & Blackburn, 2004; Valente, Etienne, & Garcia-R, 2019), it is far from unique in this regard. Human arrival and expansion in the Holocene was a major event on many other islands (Boivin et al., 2016; Louys et al., 2021), leading to extinctions, changes in community structure of plants and animals, and wholesale shifts in the structure and function of insular ecosystems (Crowley, 2010; Louys et al., 2021; G. L. Perry et al., 2014; Steadman, 1995; Wood et al., 2017).

Islands across the Pacific Ocean were populated at different times during the Polynesian expansion (Wilmshurst et al., 2011), often resulting in extreme declines in biodiversity. Among the most heavily impacted islands was Rapa Nui/Easter Island, which lost its entire endemic forest cover following the arrival of Polynesian colonists (Diamond, 2007). Similarly, Polynesians colonized the Hawaiian archipelago in the early 1200s (Rieth et al., 2011), resulting in a greater loss of native vertebrates (birds) than that following their colonization of New Zealand (Steadman, 1995). Yet each of the Pacific Islands was unique, both in their endemic biodiversity, and in their capacity to support human populations (Kirch, 1980). This surely resulted in different patterns of human population growth and spread across the archipelagos of the Pacific, and different speeds and possibly different mechanisms of biodiversity loss.

In the Indian Ocean, a similar scenario of human colonisation and extinction befell the ratite elephant birds of Madagascar (Hansford & Turvey, 2018; Hawkins & Goodman, 2003), among other species. The patterns and consequences of human colonisation of Madagascar are even more uncertain than those of New Zealand or Hawaii, with continuing debates over the latency between human colonisation and extinctions (Anderson et al., 2018; Hixon et al., 2021), along with the putative driving forces (Hansford et al., 2021). Likewise, the Caribbean islands lost many endemic vertebrates during the late Holocene (beginning around 6000 BP) (Cooke et al., 2017), however the spatiotemporal signatures and anthropogenic contribution to these extinctions remains contested (Orihuela et al., 2020).

In each of these cases, the process-explicit modelling approach we used to reconstruct island colonisation of humans across New Zealand could help untangle the potential interdependence between the dynamics of first colonists of an archipelago and the subsequent demographic, geographic and ecological dynamics of its native biota. At a minimum, this would require a dated archaeological record, climate data and ideally either an independent, direct estimate of population size following colonisation (as used here), or one inferred from molecular data (Fordham, Brook, Moritz, & Nogués-Bravo, 2014).

## Conclusions

The integration of accurately dated archaeological evidence and spatially explicit population models using a pattern-oriented paradigm enabled reliable and plausible simulations of Māori colonisation and expansion across New Zealand at a fine spatiotemporal resolution. In comparison to commonly used statistical approaches for reconstructing human migration, the modelling protocol we implemented has an advantage in that it can identify the demographic and environmental drivers of rapid colonisation events, including those that took place during periods of climatic stability, producing high resolution projections of abundance patterns that pinpoint migration routes. This is the very information needed to establish how human activities transformed island biodiversity.

Our new approach for reconstructing island colonization by humans has the potential to address outstanding questions concerning the spatiotemporal dynamics of humanity and their ecological impacts on native insular biotas of islands across the Pacific, as well as those of the Caribbean, Mediterranean, Mascarenes and Madagascar. The framework is flexible to future refinements, including the addition of different population growth models, different targets based on new archaeological and palaeobiological information, and different simulations of past climate and environmental change.

## Supporting information

Supplementary Methods and Results

## REFERENCES

Anderson, A. (1991). The chronology of colonisation in New Zealand. Antiquity, 767–795.

Anderson, A. (2016). The making of the Māori middle ages. Journal of New Zealand Studies, 23, 2–18.

Anderson, A. (2017). Changing perspectives upon Māori colonisation voyaging. Journal of the Royal Society of New Zealand, 47, 222–231.

Anderson, A., Binney, J., & Harris, A. (2015). Tangata Whenua: A History.. Wellington, New Zealand: Bridget Williams Books.

Anderson, A., Clark, G., Haberle, S., Higham, T., Nowak-Kemp, M., Prendergast, A., Camens, A. (2018). New evidence of megafaunal bone damage indicates late colonization of Madagascar. PLoS ONE, 13, p.e0204368.

Anderson, A., & Petchey, F. (2020). The transfer of kumara (Ipomoea batatas) from East to South Polynesia and its dispersal in New Zealand. Journal of the Polynesian Society, 129, 351–382.

Beyer, R. M., Krapp, M., Eriksson, A., & Manica, A. (2021). Climatic windows for human migration out of Africa in the past 300,000 years. Nature Communications, 12, 1–10.

Boivin, N. L., Zeder, M. A., Fuller, D. Q., Crowther, A., Larson, G., Erlandson, J. M., Petraglia, M. D. (2016). Ecological consequences of human niche construction: Examining long-term anthropogenic shaping of global species distributions. Proceedings of the National Academy of Sciences, 113, 6388–6396.

Boyer, A. G., & Jetz, W. (2014). Extinctions and the loss of ecological function in island bird communities. Global Ecology and Biogeography, 23, 679–688.

Brown, A. A., & Crema, E. R. (2019). Māori population growth in pre-contact New Zealand: Regional population dynamics inferred from summed probability distributions of radiocarbon dates. The Journal of Island and Coastal Archaeology, 14, 1–19.

Burney, D. A., & Flannery, T. F. (2005). Fifty millennia of catastrophic extinctions after human contact. Trends in Ecology & Evolution, 20, 395–401.

Channell, R., & Lomolino, M. V. (2000). Dynamic biogeography and conservation of endangered species. Nature, 403, 84–86.

Cooke, S. B., Dávalos, L. M., Mychajliw, A. M., Turvey, S. T., & Upham, N. S. (2017). Anthropogenic extinction dominates Holocene declines of West Indian mammals. Annual Review of Ecology, Evolution, and Systematics, 48, 301–327.

Crowley, B. E. (2010). A refined chronology of prehistoric Madagascar and the demise of the megafauna. Quaternary Science Reviews, 29, 2591–2603.

Csilléry, K., Blum, M. G. B., Gaggiotti, O. E., & François, O. (2010). Approximate Bayesian Computation (ABC) in practice. Trends in Ecology and Evolution, 25, 410–418.

Csilléry, K., François, O., & Blum, M. G. B. (2012). abc: an R package for approximate Bayesian computation (ABC). Methods in Ecology and Evolution, 3, 475–479.

Diamond, J. (1985). The riddle of the ancient mariners. The Sciences, 25, 42–46.

Diamond, J. (1997). Guns, Germs and Steel. New York: W.W. Norton & Company.

Diamond, J. (2007). Easter island revisited. Science, 317, 1692–1694.

Douglass, K., Hixon, S., Wright, H. T., Godfrey, L. R., Crowley, B. E., Manjakahery, B., Radimilahy, C. (2019)., 2019. A critical review of radiocarbon dates clarifies the human settlement of Madagascar. Quaternary Science Reviews, 221, 105878.

Duncan, R. P., & Blackburn, T. M. (2004). Extinction and endemism in the New Zealand avifauna. Global Ecology and Biogeography, 13, 509–517.

Elith, J., Leathwick, J. R., & Hastie, T. (2008). A working guide to boosted regression trees. Journal of Animal Ecology, 77, 802–813.

Ellis, E. C. (2021). Land use and ecological change: A 12,000 year history. Annual Review of Environment and Resources, 46, 1–33.

Eriksson, A., Betti, L., Friend, A. D., Lycett, S. J., Singarayer, J. S., von Cramon-Taubadel, N., Manica, A. (2012). Late Pleistocene climate change and the global expansion of anatomically modern humans. Proceedings of the National Academy of Sciences, 109, 16089.

Finlayson, C. (2005). Biogeography and evolution of the genus Homo. Trends in Ecology & Evolution, 20, 457–463.

Fordham, D. A., Brook, B. W., Moritz, C., & Nogués-Bravo, D. (2014). Better forecasts of range dynamics using genetic data. Trends in Ecology and Evolution, 29, 436–443.

Fordham, D. A., Brown, S. C., Akçakaya, H. R., Brook, B. W., Haythorne, S., Manica, A., Rahbek, C. (2022). Process-explicit models reveal pathway to extinction for woolly mammoth using pattern-oriented validation. Ecology Letters, 25, 125–137.

Fordham, D. A., Haythorne, S., & Brook, B. W. (2016). Sensitivity Analysis of Range Dynamics Models (SARDM): Quantifying the influence of parameter uncertainty on forecasts of extinction risk from global change. Environmental Modelling & Software, 83, 193–197.

Fordham, D. A., Haythorne, S., Brown, S. C., Buettel, J. C., & Brook, B. W. (2021). poems: R package for simulating species’ range dynamics using pattern-oriented validation. Methods in Ecology and Evolution, 12, 2364–2371.

Fordham, D. A., Jackson, S. T., Brown, S. C., Huntley, B., Brook, B. W., Dahl-Jensen, D., Wilmshurst, J. M. (2020). Using paleo-archives to safeguard biodiversity under climate change. Science, 369, eabc5654.

Fordham, D. A., Saltré, F., Haythorne, S., Wigley, T. M., Otto-Bliesner, B. L., Chan, K. C., & Brook, B. W. (2017). PaleoView: a tool for generating continuous climate projections spanning the last 21 000 years at regional and global scales. Ecography, 40, 1348–1358.

Furey, L. (2006). Maori Gardening: An Archaeological Perspective. Wellington: Department of Conservation.

Gelman, A., Hwang, J., & Vehtari, A. (2014). Understanding predictive information criteria for Bayesian models. Statistics and Computing, 24, 997–1016.

Giampoudakis, K., Marske, K. A., Borregaard, M. K., Ugan, A., Singarayer, J. S., Valdes, P. J., Nogués-Bravo, D. (2017). Niche dynamics of Palaeolithic modern humans during the settlement of the Palaearctic. Global Ecology and Biogeography, 26, 359–370.

Goldberg, A., Mychajliw, A. M., & Hadly, E. A. (2016). Post-invasion demography of prehistoric humans in South America. Nature, 532, 232–235.

Greig, K., & Rawlence, N. J. (2021). The potential contribution of kurī (Polynesian dog) to the ecological impacts of the human settlement of Aotearoa New Zealand. EcoEvoRxiv, doi:10.32942/osf.io/khfg8.

Grimm, V., & Railsback, S. F. (2012). Pattern-oriented modelling: a ‘multi-scope’ for predictive systems ecology. Philosophical Transactions of the Royal Society B, 367, 298–310.

Hamel, J. (1982). South Otago. In N. Prickett (Ed.), The First Thousand Years: Regional Perspectives in New Zealand Archaeology (pp. 129–150). Palmerston North: Dunmore Press.

Hansford, J. P., Lister, A. M., Weston, E. M., & Turvey, S. T. (2021). Simultaneous extinction of Madagascar’s megaherbivores correlates with late Holocene human-caused landscape transformation. Quaternary Science Reviews, 263, 106996.

Hansford, J. P., & Turvey, S. T. (2018). Unexpected diversity within the extinct elephant birds (Aves: Aepyornithidae) and a new identity for the world’s largest bird. Royal Society Open Science, 5, 181295.

Hawkins, A. F. A., & Goodman, S. M. (2003). Introduction to the birds. In S. M. Goodman & J. P. Benstead (Eds.), The Natural History of Madagascar (pp. 1019–1044). Chicago, USA: University of Chicago Press.

Haythorne, S., Fordham, D. A., Brown, S. C., Buettel, J. C., & Brook, B. W. (2021). poems: pattern-oriented ensemble modeling system. The Comprehensive R Archive Network.

Hixon, S. W., Douglass, K. G., Crowley, B. E., Rakotozafy, L. M. A., Clark, G., Anderson, A., Mbola, B. (2021). Late Holocene spread of pastoralism coincides with endemic megafaunal extinction on Madagascar. Proceedings of the Royal Society B, 20211204.

Holdaway, R. N., Allentoft, M. E., Jacomb, C., Oskam, C. L., Beavan, N. R., & Bunce, M. (2014). An extremely low-density human population exterminated New Zealand moa. Nature Communications, 5, 5436.

Holdaway, R. N., & Jacomb, C. (2000). Rapid extinction of the moas (Aves: Dinornithiformes): model, test, and implications. Science, 287, 2250–2254.

Holdaway, S. J., Emmitt, J., Furey, L., Jorgensen, A., O’Regan, G., Phillipps, R., Ladefoged, T. N. (2019). Māori settlement of New Zealand: The Anthropocene as a process. Archaeology in Oceania, 54, 17–34.

Honkaniemi, J., Rammer, W., & Seidl, R. (2021). From mycelia to mastodons A general approach for simulating biotic disturbances in forest ecosystems. Environmental Modelling & Software, 138, 104977.

Kinaston, R. L., Walter, R. K., Jacomb, C., Brooks, E., Tayles, N., Halcrow, S. E., Shaw, B. (2013). The first New Zealanders: patterns of diet and mobility revealed through isotope analysis. PLoS ONE, 8, p.e64580.

Kirch, P. V. (1980). Polynesian prehistory: Cultural adaptation in island ecosystems. American Scientist, 68, 39–48.

Locatelli, E., Due, R. A., van den Bergh, G. D., & van den Hoek Ostende, L. W. (2012). Pleistocene survivors and Holocene extinctions: the giant rats from Liang Bua (Flores, Indonesia). Quaternary International, 281, 47–57.

Louys, J., Braje, T. J., Chang, C. H., Cosgrove, R., Fitzpatrick, S. M., Fujita, M., O’Connor, S. (2021). No evidence for widespread island extinctions after Pleistocene hominin arrival. Proceedings of the National Academy of Sciences, 118, e2023005118.

McGlone, M., Anderson, A., & Holdaway, R. (1994). An ecological approach to the Polynesian settlement of New Zealand. In D. Sutton (Ed.), The Origin of the First New Zealanders (pp. 136–163). Auckland: Auckland University Press.

McGlone, M. S., & Wilmshurst, J. M. (1999). Dating initial Maori environmental impact in New Zealand. Quaternary International, 59, 5–16.

Mellars, P. (2006). Why did modern human populations disperse from Africa ca. 60,000 years ago? A new model. Proceedings of the National Academy of Sciences, 103, 9381–9386.

Mellin, C., Russell, B. D., Connell, S. D., Brook, B. W., & Fordham, D. A. (2012). Geographic range determinants of two commercially important marine molluscs. Diversity and Distributions, 18, 133–146.

Nielsen, R., Akey, J. M., Jakobsson, M., Pritchard, J. K., Tishkoff, S., & Willerslev, E. (2017). Tracing the peopling of the world through genomics. Nature, 541, 302–310.

Nogué, S., Santos, A. M., Birks, H. J. B., Björck, S., Castilla-Beltrán, A., Connor, S., Steinbauer, M. J. (2021). The human dimension of biodiversity changes on islands. Science, 372, 488–491.

Orihuela, J., Viñola, L. W., Vázquez, O. J., Mychajliw, A. M., de Lara, O. H., Lorenzo, L., & Soto-Centeno, J. A. (2020). Assessing the role of humans in Greater Antillean land vertebrate extinctions: New insights from Cuba. Quaternary Science Reviews, 249, 106597.

Perry, G. L., Wainwright, J., Etherington, T. R., & Wilmshurst, J. M. (2016). Experimental simulation: using generative modeling and palaeoecological data to understand human-environment interactions. Frontiers in Ecology and Evolution, 4, 109.

Perry, G. L., Wheeler, A. B., Wood, J. R., & Wilmshurst, J. M. (2014). A high-precision chronology for the rapid extinction of New Zealand moa (Aves, Dinornithiformes). Quaternary Science Reviews, 105, 126–135.

Perry, G. L. W., Wilmshurst, J. M., McGlone, M. S., & Napier, A. (2012). Reconstructing spatial vulnerability to forest loss by fire in pre-historic New Zealand. Global Ecology and Biogeography, 21, 1029–1041.

Pilowsky, J. A., Colwell, R. K., Rahbek, C., & Fordham, D. A. (2022). Process-explicit models reveal the structure and dynamics of biodiversity. Science Advances, doi: 10.1126/sciadv.abj2271.

Pool, D. I. (1991). Te iwi Maori: a New Zealand Population, Past, Present and Projected. Auckland, New Zealand: Auckland University Press.

Potts, J. M., & Elith, J. (2006). Comparing species abundance models. Ecological Modelling, 199, 153–163.

R Core Team. (2021). R: A language and environment for statistical computing. Vienna, Austria: R Foundation for Statistical Computing.

Rangel, T. F., Edwards, N. R., Holden, P. B., Diniz-Filho, J. A. F., Gosling, W. D., Coelho, M. T. P., Colwell, R. K. (2018). Modeling the ecology and evolution of biodiversity: Biogeographical cradles, museums, and graves. Science, 361, 6399.

Rawlence, N. J., Perry, G. L. W., Smith, I. W. G., Scofield, R. P., Tennyson, A. J. D., Matisoo-Smith, E. A., Waters, J. M. (2015). Radiocarbon-dating and ancient DNA reveal rapid replacement of extinct prehistoric penguins. Quaternary Science Reviews, 112, 59–65.

Ridout, M., Demétrio, C. G., & Hinde, J. (1998). Models for count data with many zeros. Proceedings of the XIXth International Biometric Conference 19, 179–192.

Rieth, T. M., Hunt, T. L., Lipo, C., & Wilmshurst, J. M. (2011). The 13th century Polynesian colonization of Hawai’i Island. Journal of Archaeological Science, 38, 2740–2749.

Russell, J. C., & Kueffer, C. (2019). Island biodiversity in the Anthropocene. Annual Review of Environment and Resources, 44, 31–60.

Saltelli, A., Tarantola, S., & Campolongo, F. (2000). Sensitivity analysis as an ingredient of modeling. Statistical Science, 15, 377–395.

Smith, I. (2013). Pre-European Maori exploitation of marine resources in two New Zealand case study areas: species range and temporal change. Journal of the Royal Society of New Zealand, 43, 1–37.

Steadman, D. W. (1995). Prehistoric extinctions of Pacific island birds: Biodiversity meets zooarchaeology. Science, 267, 1123–1131.

Timmermann, A., & Friedrich, T. (2016). Late Pleistocene climate drivers of early human migration. Nature, 538, 92–95.

Valente, L., Etienne, R. S., & Garcia-R, J. C. (2019). Deep macroevolutionary impact of humans on New Zealand’s unique avifauna. Current Biology, 29, 2563–2569.

Wallace, A. R. (1876). The Geographical Distribution of Animals; With a Study of the Relations of Living and Extinct Faunas as Elucidating the Past Changes of the Earth’s Surface. New York: Harper and Brothers.

Walter, R., Buckley, H., Jacomb, C., & Matisoo-Smith, E. (2017). Mass migration and the Polynesian settlement of New Zealand. Journal of World Prehistory, 30, 351–376.

Wanner, H., Beer, J., Butikofer, J., Crowley, T. J., Cubasch, U., Flückiger, J., Widmann, M. (2008). Mid- to Late Holocene climate change: An overview. Quaternary Science Reviews, 27, 1791–1828.

Whyte, A. L., Marshall, S. J., & Chambers, G. K. (2005). Human evolution in Polynesia. Human Biology, 77, 157–177.

Wiegand, T., Moloney, K. A., Naves, J., & Knauer, F. (1999). Finding the missing link between landscape structure and population dynamics: A spatially explicit perspective. The American Naturalist, 154, 605–627.

Wilmshurst, J. M., Anderson, A. J., Higham, T. F., & Worthy, T. H. (2008). Dating the late prehistoric dispersal of Polynesians to New Zealand using the commensal Pacific rat. Proceedings of the National Academy of Sciences, 105, 7676–7680.

Wilmshurst, J. M., Higham, T. F., Allen, H., Johns, D., & Phillips, C. (2004). Early Maori settlement impacts in northern coastal Taranaki, New Zealand. New Zealand Journal of Ecology, 28, 167–179.

Wilmshurst, J. M., Hunt, T. L., Lipo, C. P., & Anderson, A. (2011). High-precision radiocarbon dating shows recent and rapid initial human colonization of East Polynesia. Proceedings of the National Acadamy of Science, 108, 1815–1820.

Wood, J. R. (2008). Moa (Aves: Dinornithiformes) nesting material from rockshelters in the semi-arid interior of South Island, New Zealand. Journal of the Royal Society of New Zealand, 38, 115–129.

Wood, J. R., Alcover, J. A., Blackburn, T. M., Bover, P., Duncan, R. P., Hume, J. P., Wilmshurst, J. M. (2017). Island extinctions: processes, patterns, and potential for ecosystem restoration. Environmental Conservation, 44, 348–358.

Zhu, D., Galbraith, E. D., Reyes-García, V., & Ciais, P. (2021). Global hunter-gatherer population densities constrained by influence of seasonality on diet composition. Nature Ecology & Evolution, 5, 1536–1545.

