## Supplementary Methods and Results for "Reconstructing colonization dynamics to establish how human activities transformed island biodiversity"

**SUPPORTING METHODS**

Archaeological data

Training data for the human density model came from the New Zealand Radiocarbon database (https://www.waikato.ac.nz/nzcd/index.html) (n = 1671). The quality of the dates was assessed using established criteria (Wilmshurst, Anderson, Higham, & Worthy, 2008), resulting in 645 reliably dated archaeological records, which we supplemented with 55 further reliable records (Brown & Crema, 2019).

The New Zealand Radiocarbon database ends at 2003 and does not record archaeological sites that have not been ^14^C dated. Therefore, we also accessed all archaeological records maintained by the New Zealand Archaeological Association (http://www.archsite.org.nz/Default.aspx). The archaeological records were filtered using age categories for each site. Only sites identified as pre-1769 and unequivocally relating to traditional Māori activities were retained. This meant removing administrative, botanical, commercial, flax and flour milling, healthcare, historic, industrial, military, mining, missionary and religious, shipwreck, timber milling, whaling station and unclassified records and locations. Because the construction of fortifications is associated with Māori society long after the putative settlement and colonisation period (Allen, 2016; Pearce & Pearce, 2010), we removed all sites associated with *pa*. In total 21, 644 recorded archaeological sites were retained for comparison with the ^14^C artefact data set.

To assess whether the smaller, but more reliable, ^14^C database was representative of the broader spatial pattern of Māori occupation during the early Polynesian colonisation period, we quantified the degree of spatial correlation between the filtered New Zealand Archaeological Association data and our ^14^C database. To do this, New Zealand was divided into grid cells of approximately 10 km^2^, 25 km^2^ and 50 km^2^ resolution, and the numbers of ^14^C dated sites and undated archaeological sites within each grid cell were summed. We quantified spatial associations in the count data of both dated and undated sites using spatial analysis by distance indices (SADIE; (J. N. Perry, 1995, 1998; J. N. Perry & Hewitt, 1991); Supplementary Figure 1) and a correlation association index (Winder et al., 2019) using the *‘epiphy’* v 0.3.4 R package at the three separate spatial resolutions.

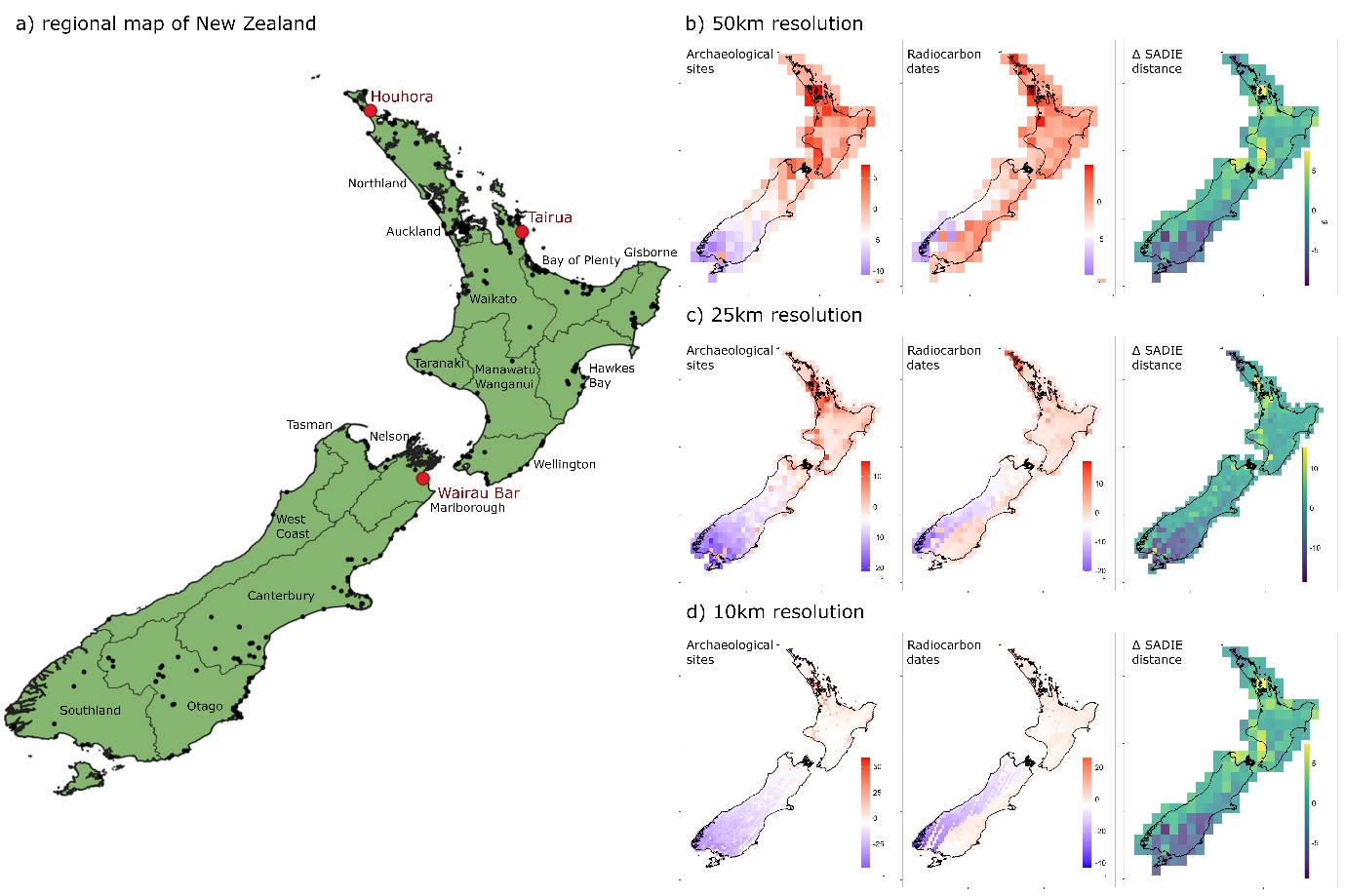

**Supplementary Figure 1:** New Zealand, showing **a)** regions of modern New Zealand, the location of radiocarbon-dated evidence of Māori occupation (black) and the oldest known sites of Māori settlement (red). These resulted in spatial correlations of evidence of Māori occupation at **b)** 50km^2^ resolution, **c)** 25km^2^ resolution and **d)** 10km^2^ resolution. The red-blue plots for the number of sites containing archaeological evidence of Māori occupation, and ^14^C dated artefacts of Māori occupation indicate the relative distance that each grid cell is from a uniform distribution, estimated by spatial analysis by distance indices (SADIE). The cell-by-cell differences in SADIE patterns between the artefacts and the archaeological sites indicated that, while the magnitude of the differences is lowest at 50 km^2^ resolution, the extreme differences encompass greater land area, while at the 10 km^2^ resolution, the magnitude of differences was substantial. We found the greatest concordance between artefacts and archaeological site records occurred at the 25km^2^ resolution.

Climate data

The influence of climate on the abundance of Māori archaeological sites (a proxy for relative abundance; (Goldberg, Mychajliw, & Hadly, 2016)) was modelled using annual average rainfall (mL) and the average monthly temperature (°C) for the coldest and warmest three-month periods. These climatic factors have previously been used in statistical models of human migration (Giampoudakis et al., 2017). We used PaleoView v1.5.1 (Fordham et al., 2017) to generate climate projections for the period of Māori colonisation. To do this we calculated 30 year average anomalies at 25 year intervals from a baseline of 1975 (1960-1989) for the period from 935 CE to 1970 CE. To capture important orthographic elements in New Zealand’s climate, these anomalies were downscaled to a 0.30° x 0.30° grid cell resolution using a change factor method (Fordham, Wigley, & Brook, 2011) and climate data from the New Zealand National Institute of Water and Atmospheric Research (NIWA) and the University of East Anglia Climatic Research Unit (CRU) (Harris, Osborn, Jones, & Lister, 2020). We could not downscale the PaleoView data directly to the NIWA data because PaleoView provides 30-year average climate data up until 1975, while NIWA have 30-year average data from 1985. To address this temporal mismatch, we first adjusted the NIWA climate data for observed climatic changes in New Zealand between 1975 and 1985 using data from CRU. This harmonisation process has been used elsewhere in a similar way (Fordham, Brown, et al., 2021).

Environmental predictors

The influence of environmental context on Māori archaeological sites was modelled using net primary productivity (NPP), evapotranspiration, distance to navigable rivers and the coastline, and the percentage of each 25 km^2^ grid cell that was steeper than 20 degrees slope using a 1 arc-second resolution digital elevation model (Landcare Research New L. R. N. Zealand, 2010). High net primary productivity (NPP) is associated with high population densities of pre-agricultural human populations up to a threshold, beyond which human population density becomes largely independent of NPP (Tallavaara, Eronen, & Luoto, 2018). The positive relationship between NPP and population density of pre-agricultural human populations is hypothesised to relate to increased resource availability (Binford, 2001; Tallavaara et al., 2018). Using monthly minimum temperature and monthly maximum temperature (described above), we calculated NPP using the empirical Miami model (Lieth, 1973), which can produce robust estimates of current-day NPP (Adams, White, & Lenton, 2004):

$NPP=min\binom{\frac{3000}{1+ e^{1.315-0.199 \times T_{a}}}}{3000 \times(1-e^{-0.000644 \times P})}$	(1)

Where *T_a_* is temperature and *P* is precipitation.

Evapotranspiration was used as a proxy for the capacity for horticultural plant growth under the specific climatic conditions in each cell (Droogers & Allen, 2002):

${ET}_{0}=0.0013 \times\left( 0.408 \times R_{s} \right)\times\left( T_{a}+17.0 \right)\times{(\left( T_{max}-T_{min} \right) -0.0123\times P)}^{0.76}$	(2)

Where *R_s_* is solar radiation, *T_a_* is the average annual temperature, *P* is precipitation, *T_min_* is the average minimum temperature and *T_max_* is the average maximum temperature.

We applied an upper slope of 20 degrees as the limit to Māori habitat suibtaility because Māori gardening generally required gentle slopes, and has rarely been recorded on slopes > 15-20 degrees (Allen, 2016; Furey, 2006). We quantified distance to navigable rivers and distance to the coastline. We identified navigable rivers as those with annual flow of >10m.s^-1^ from a data set collated by Booker (2015), with the assumption that these flow rates tend to be associated with the larger river valleys that could provide access from the coast to the interior, even in seasons when water depth didn’t allow navigation. We also calculated distance from all navigable waters (coast and rivers combined).

Hurdle modelling

To estimate spatial patterns of Māori relative abundance we built a decomposed hurdle model, allowing us to train the occurrence and abundance steps of the model on different environmental factors (Potts & Elith, 2006; Ridout, Demétrio, & Hinde, 1998), conducting model diagnostics at each step (Mellin, Russell, Connell, Brook, & Fordham, 2012). We modelled the probability of presence using a binomial distribution with the ‘*gbm’* v2.1.8 package (Greenwell, Boehmke, J., & Developers, 2019) and functions provided by Elith, Leathwick, and Hastie (2008). The 155 cells with dated archaeological sites were classified as binary “presence” locations. The remaining cells were used as “absence”, given that they roughly equated to the number of presence locations (Elith et al., 2008). This allowed us to identify the four variables that most strongly contributed to patterns in ^14^C dated Māori presence (Supplementary Figure 2). The second step of the hurdle model consisted of fitting a model of site abundance, conditional on modelled presence (i.e., only in regions of New Zealand where the likelihood of occurrence was above the TSS threshold). The model of Māori archaeological site abundance was initially fitted to all the climatic and environmental variables and then simplified to identify the four variables most strongly driving the patterns of site abundance (Supplementary Figure 2). To capture uncertainty in model fits we bootstrapped the data. Specifically, we took 5000 replicate subsamples, retaining 70% of the ^14^C dated Māori artefact data for training. Each replicate model was projected across New Zealand to generate a relative density estimate of Māori. The mean and standard deviation of these 5000 replicate estimates were retained as the Māori relative density templates for spatially explicit population modelling.

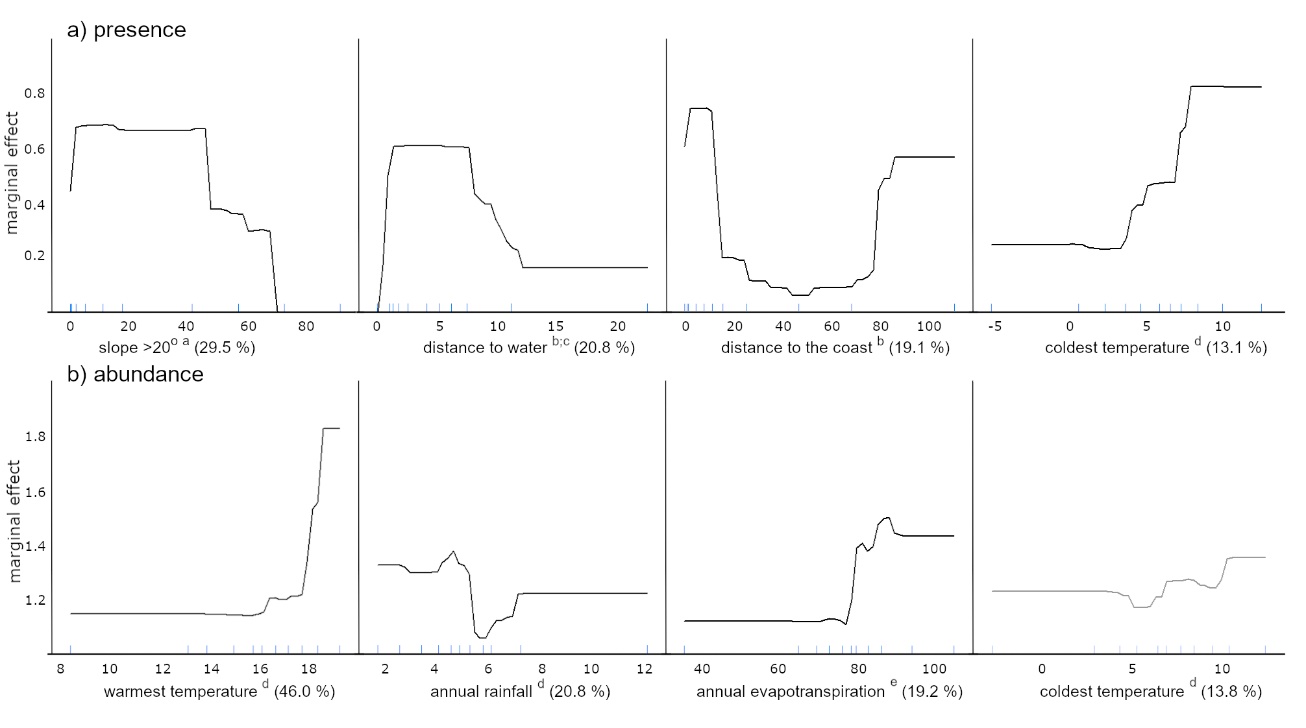

**Supplementary Figure 2:** Climatic and environmental correlates of the presence and abundance of Māori archaeological sites. Percentage values in the two columns indicate the contribution that each variable made to models of presence and abundance, separately. Note that net primary productivity and distance to navigable rivers were not retained in either model. Superscript letters indicate data sources: a = Landcare Research New L. R. N. Zealand (2010); b= Land Information New L. I. N. Zealand (2018); c = Booker (2015); d = Fordham et al. (2017); e = Droogers and Allen (2002).

Probability of presence of Māori settlement was estimated for all of New Zealand and used as proxy for habitat suitability (Hirzel & Le Lay, 2008). We used the True Skill Score (TSS) (Liu, Berry, Dawson, & Pearson, 2005) to calculate a threshold for applying a presence/absence cut-off to the continuous projections of probability of occurrence. We did 1000-iteration bootstraps, where we calculated TSS based on 10% of hold-out data, at 1% thresholds of occurrence between 0 and 1 using the *‘TSS’* v0.3.0 package (Krug, 2014).

The accuracy of the presence model was assessed using independent data as well as cross validation. The presence and absence of archaeological sites in the New Zealand Archaeological Association database was used as an “out of sample” validation. The Boyce Index was used to validate model projections (Boyce, Vernier, S.E., & Schmiegelow, 2002; Hirzel, Le Lay, Helfer, Randin, & Guisan, 2006) using the *‘ecospat’* v3.2.1 package (function ‘ecospat.boyce; (Di Cola et al., 2017)). The abundance predictions of our model were compared against the spatial patterns of archaeological site density from the New Zealand Archaeological Association database as an out of sample validation using SADIE; (J. N. Perry, 1995, 1998; J. N. Perry & Hewitt, 1991) and the *‘epiphy’* v0.3.4 package (Gigot, 2018).

SEPM Landscape suitability

In order to ensure sufficient (and realistic) separation of the North and South Islands in model simulations, the relative abundance (and its SD) from the hurdle model was downscaled to 20 km^2^ resolution using bilinear interpolation. We then generated thousands of potential landscapes of relative abundance for Māori by resampling from the rasters of mean Māori relative abundance and its standard deviation. To ensure that grid cells in close spatial association were more similar than spatially distant grid cells, we resampled these landscapes of Māori relative abundance using an estimate of environmental correlation (Gregory et al., 2014), which we describe below.

Population Growth

We reconstructed population size at annual intervals for the time period 1230 – 1850 AD using a broad range of population growth rates and founding populations and an exponential population growth function (Holdaway et al., 2014; Holdaway & Jacomb, 2000). We assumed a single founding population, ranging in size between 200, 300, 400, 500 and 600 individuals, growing at annual rates of 0.5, 1, 1.3, 1.4, 1.5 and 2.2%. With a population growth window of approximately 600 years, between 1280 and 1769 (Pool, 1991; Wilmshurst et al., 2008), population size in 1769 ranged from 98,092 – 146,385, suggesting that the range of model parameters provided a suitably large but plausible band of growth estimates for reconstructing Māori colonisation, expansion and growth. The population growth model that we applied was characterised with 30 different plausible parameterisations of a two-parameter exponential model (Supplementary Figure 3) (Holdaway et al., 2014; Holdaway & Jacomb, 2000):

$N_{t}=R^{time} \times N_{0}$	(3)

Where *R* is the intrinsic rate of population growth, *N_0_* is the founding population, and *time* is the number of years since the population was founded.

We chose not to use a logistic growth function (Brown & Crema, 2019) because it simulated overly rapid rates of population growth early in the time series (exceeded the growth rates generally accepted for subsistence human population (Bentley, 1985; Walker et al., 2006)), and required a colonisation date well before 1230 and 1314 AD (G. L. Perry, Wheeler, Wood, & Wilmshurst, 2014; Wilmshurst et al., 2008) to simulate the target population size in 1769.

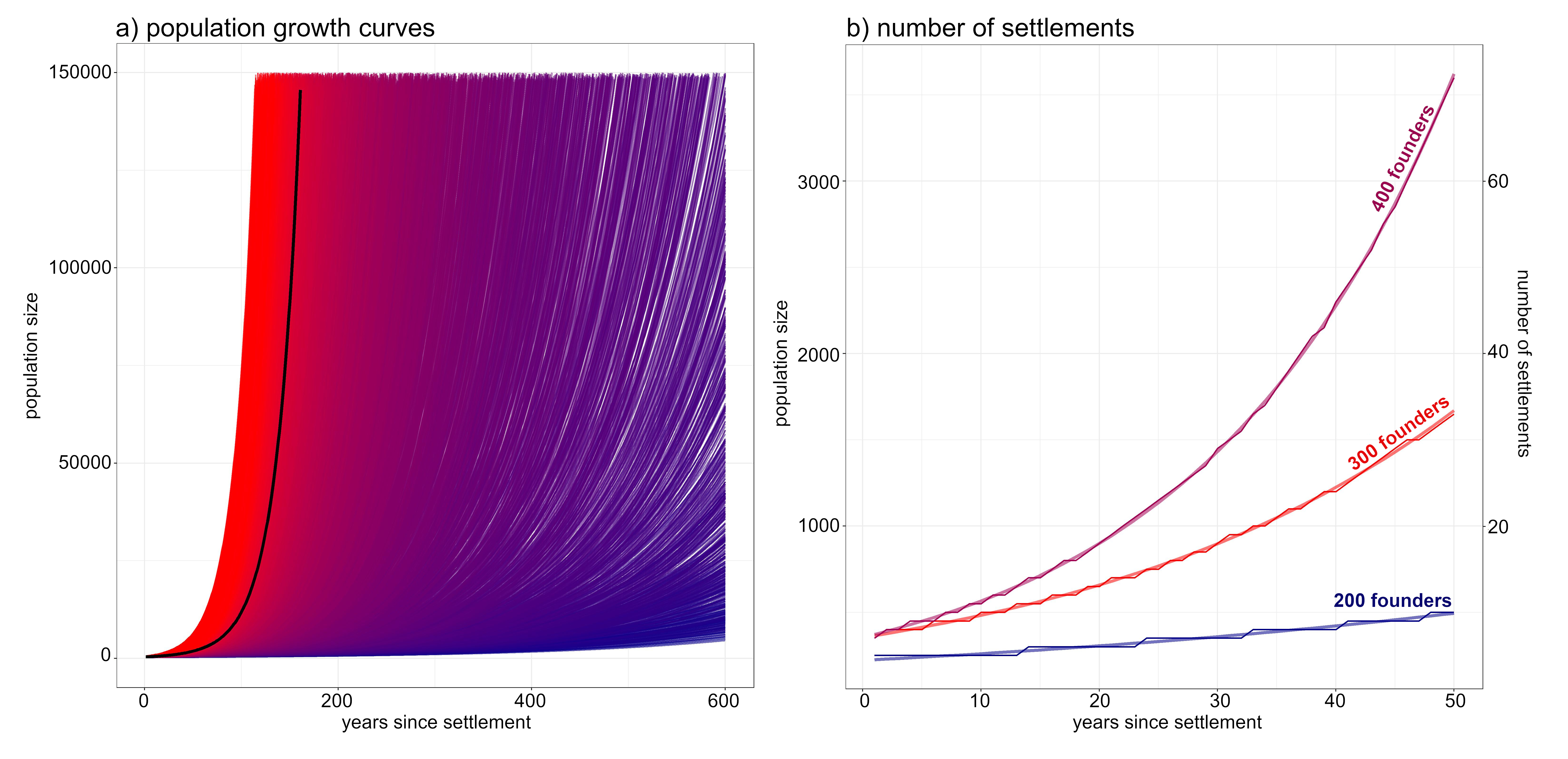

**Supplementary Figure 3:** a) Population growth curves for Māori. Curves show output of the full range of plausible models of Māori population growth (Holdaway et al., 2014; Holdaway & Jacomb, 2000). The heavy black line indicates the multi-model average. b) Settlement numbers resulting from different growth curves. Each curve is identical, but for the founding population size, but the number of populations of a minimum size (in this case 50 individuals) increases stepwise at different rates according to different population growth rates.

*Spatially explicit population model*

We modelled range expansion as a function of population size and habitable neighbourhoods using a lattice-type spatially explicit population model (SEPM) coded in the R package *‘poems’* v1.0.1 (Haythorne, Fordham, Brown, Buettel, & Brook, 2021). A user-defined function aggregated grid cells into spatial neighbourhoods with foraging radii ranging from 30-70 km. At each time step, neighbourhoods available for colonisation were identified using the spatio-temporal estimate of landscape abundance. Neighbourhoods with highest potential relative abundances were colonised first.

We modelled the movement of communities of people rather than individuals. We did this by varying minimum community size between 20-80 people. As such, we simulated dispersal of the Māori population across New Zealand by first populating the areas with the conditions most suitable for high relative density of archaeological sites, then dispersing into areas of relatively lower suitability as the total population grew (Figure 3). While a key assumption in the model was that Māori were able to move long distances over short periods of time, including by boat (Irwin et al., 2017; Walter, Buckley, Jacomb, & Matisoo-Smith, 2017), there was a tendency to move shorter distances early in the simulation owing to highly suitable areas for colonisation being spatially autocorrelated and non-random sampling.

We used spatial correlation of climatic conditions to guide the colonisation of neighbourhoods, given that human migration was strongly driven by climate (Timmermann & Friedrich, 2016). Previously, this approach has been used to simulate spatial correlation in demographic processes for vertebrates (Fordham et al., 2013; Fordham et al., 2018). To do this we analysed monthly NIWA precipitation data for the period 1961 to 1985 from 52 meteorological stations across New Zealand (https://niwa.co.nz/education-and-training/schools/resources/climate) as a proxy for environmental correlation (Fordham et al., 2012). We measured spatio-temporal environmental correlation separately for the North and South Islands using a non-centred (spline) spatial cross-correlogram implemented in the R package *‘ncf’* v1.8.1 (Bjornstad & Falck, 2001). We ran 500 bootstrap iterations to estimate 95% error bounds in the spatial correlation of temperature and precipitation. Using non-linear regression in the R package *‘nlstools’* v2.0.0 (Baty et al., 2015) we parameterised a negative exponential correlation function (Akçakaya, 1998) incorporating the patterns of correlation by distance across both islands. The resulting parameters were used to define spatial correlation in the *poems* modelling architecture.

Pattern oriented modelling

We validated our SEPM using pattern-oriented modelling (POM) methods (Fordham, Haythorne, Brown, Buettel, & Brook, 2021). To do this we varied five parameters across large but plausible ranges (Table 1) using a robust coverage of multi-dimensional parameter space (Fordham, Haythorne, & Brook, 2016). These parameters were the number of Polynesian colonists that first arrived in New Zealand (human_founding_population), the rate at which the Māori population increased following arrival in New Zealand (human_growth_rate), the year (AD) in which Polynesians colonised New Zealand (human_colonisation_time), the foraging radius (km) of each Māori community (human_foraging_distance), and the minimum number of people required to seed a new Māori settlement (human_min_neighbourhood). Parameters were varied using Latin hypercube sampling, which assigns a plausible range for each parameter and samples all subsets of its distribution once, based on subdivisions of equal probability density (Norton, 2015), resulting in a stratified random subset of parameter input values. Since there were no best estimates for our varied parameters, we used uniform sampling distributions (Fordham, Brown, et al., 2021), generating a uniform set of uninformed prior parameter estimates. This procedure produced 25,000 conceivable models with different combinations of variable parameters, each of which we ran for a single replicate (Prowse et al., 2016). Each model produced grided spatio-temporal simulations of Māori abundance, characterised by time of colonisation and subsequent population sizes across New Zealand.

Following the POM paradigm, Approximate Bayesian Computation (ABC; (Csilléry, Blum, Gaggiotti, & François, 2010)) was used to compare simulation outputs with validation targets: the percentage deviation from an estimated population size of 100, 000 - 150, 000 in 1769, and the percentage of archaeological sites that were modelled to be occupied prior to the oldest ^14^C date available there. We selected the best 1% of simulations that did best at matching the multi-variate target based on the ‘rejection’ algorithm in the R package *‘abc’* v2.1 (Csilléry, François, & Blum, 2012). We used the rejection algorithm to construct the posterior distributions of parameters because this algorithm maintains the parameter estimates as a subsample of the priors, rather than correcting them to the summary statistics of the prior distribution (Csilléry et al., 2012).

We compared differences between posterior and prior distributions of parameters by calculating the Bayes Factor (Makowski, Ben-Shachar, & Lüdecke, 2019). A non-significant Bayes Factor indicates convergence between the distribution of the parameters comprising the selected simulations (prior parameters distribution), and the pool of parameters from which they were drawn (posterior parameter distribution) (Makowski et al., 2019). Using this metric, the parameter range identified by the ABC as most accurately matching the validation targets was repeatedly resampled to constrain the plausible parameter space. Resampling was achieved by fitting a candidate set of probability density functions (normal, lognormal, Poisson, binomial, negative binomial and uniform) to each parameter and selecting the most appropriate distribution by Akaike’s Information Criterion, AIC (Burnham & Anderson, 2002). The resampled data set was assembled by drawing 25,000 new combinations of parameters to match the distributions of the model posteriors of the previous round of simulations. These models were then simulated and assessed by ABC until the Bayes Factor indicated that the selected models were statistically indistinguishable from the sample data (i.e. the Bayes Factor for each parameter was zero). This required two resampling rounds to achieve (Supplementary Table 2), however we also tested the third round of samples which were drawn for posterior predictive testing. We did posterior predictive checks (Crespi & Boscardin, 2009) to determine whether the posterior distributions result in good resemblance between simulated and observed data (Gelman, Hwang, & Vehtari, 2014). Essentially, these checks were used to identify differences in the parameter space between the sample data and the simulations selected by ABC.

SEPM Sensitivity Analyses

We did a sensitivity analysis to determine the sensitivity of the results to two model-based structural assumptions (Saltelli, Tarantola, & Campolongo, 2000): the form of the population growth model and the number of independent colonisation events. For the first, we tested the application of a logistic population growth model advocated by Brown and Crema (2019) in place of the exponential model that we used. To test this, we generated 25,000 simulations using a reparametrized human density algorithm using the growth function:

$\frac{k}{1+ \left( \frac{k- p_{0}}{p_{0}} \right)\times e^{-r \times\mathrm{time}}}$	(4)

Where time is the number of years since the colonisation year, *p_0_* is the founding population size identified at iteration two, *r* is the growth rate identified at iteration two, and *k* is the human carrying capacity, uniformly sampled between 5300 and 265,000 (Brown and Crema, 2019).

The second of structural assumption that we tested was that of a single arrival time. We constructed three alternative arrival scenarios, where the founding population arrived in two annual contingents, five annual contingents and ten annual contingents respectively. For each scenario we generated 25,000 simulations. All other components remained unchanged from the baseline SEPM.

**SUPPORTING RESULTS**

Archaeological data

The 25km^2^ resolution captured the finest degree of spatial patterning possible in the dated sites, without differing greatly from the patterns in the larger set of undated sites (Supplementary Figure 1). On this basis, all subsequent spatial modelling was undertaken at 25km^2^ resolution.

**Supplementary Table 1:** Independent tests of the Māori relative density model, tested using the area under the receiver operator curve (AUC), the Boyce Index (reported as a Spearman correlation, and the True Skill Score (TSS) for maximising the relationship between predicted probability of occurrence and presence on the landscape. Mean error and Root Mean Square Error (RMSE) of predictions against bootstrapped cross-validation.

| **Validation parameter** | **Test value** |
| --- | --- |
| AUC | 0.860 |
| Spearman Correlation | 0.956 |
| TSS | 0.660 |
| Mean error | 0.220 |
| RMSE | 5.130 |

Hurdle modelling

The hurdle models proved robust when tested using the area under the receiver operator curve (AUC), Boyce Index (reported as a Spearman correlation) and TSS (Supplementary Table 1). 10-fold cross-validation returned a mean error with respect to the likelihood of Māori settlement of 0.22, a RMSE of 5.13, from an average abundance of 4.47 ± 0.81(SD). The spatial patterns in abundance of archaeological finds remained relatively consistent between SADIE estimates trained on the archaeological site locations and on our modelled output (Supplementary Figure 4).

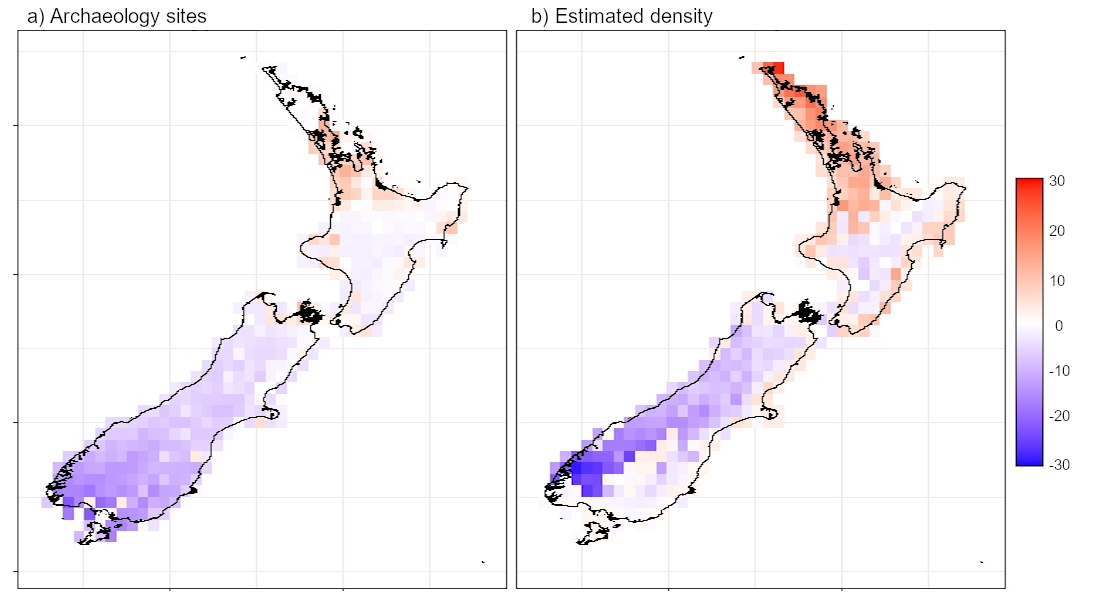

**Supplementary Figure 4:** Red-blue plots at 25 km^2^ resolution, indicating the relative distance that each grid cell is from a uniform distribution, estimated by spatial analysis by distance indices (SADIE) for a) the independent undated archaeological sites of Māori populations, and b) the population density estimates produced by our models. Both estimates agree that the North Island supported the highest positive deviation in observations, particularly along the Northland Peninsular.

Population Growth Rates

When appropriately integrated with the SEPM, growth rates at the population level (i.e. at each neighbourhood) were spatially heterogenous (Supplementary Figure 5). This spatial pattern emerged as a result of the human dispersal model preferentially growing different populations in proportion to the relative population template. When summarised by bioregions, the population growth rates generated by our model were similar to those identified by Brown and Crema (2019), with highest rates of growth on the North Island, however our model suggests that maximal growth rates were highest in areas analogous to the Central economic subregion defined by Brown and Crema (2019) (approximately 1.013), followed by the Central economic subregion (approximately 1.012). The Southern economic subregion had much lower growth rates (approximately 1.01; Supplementary Figure 5)

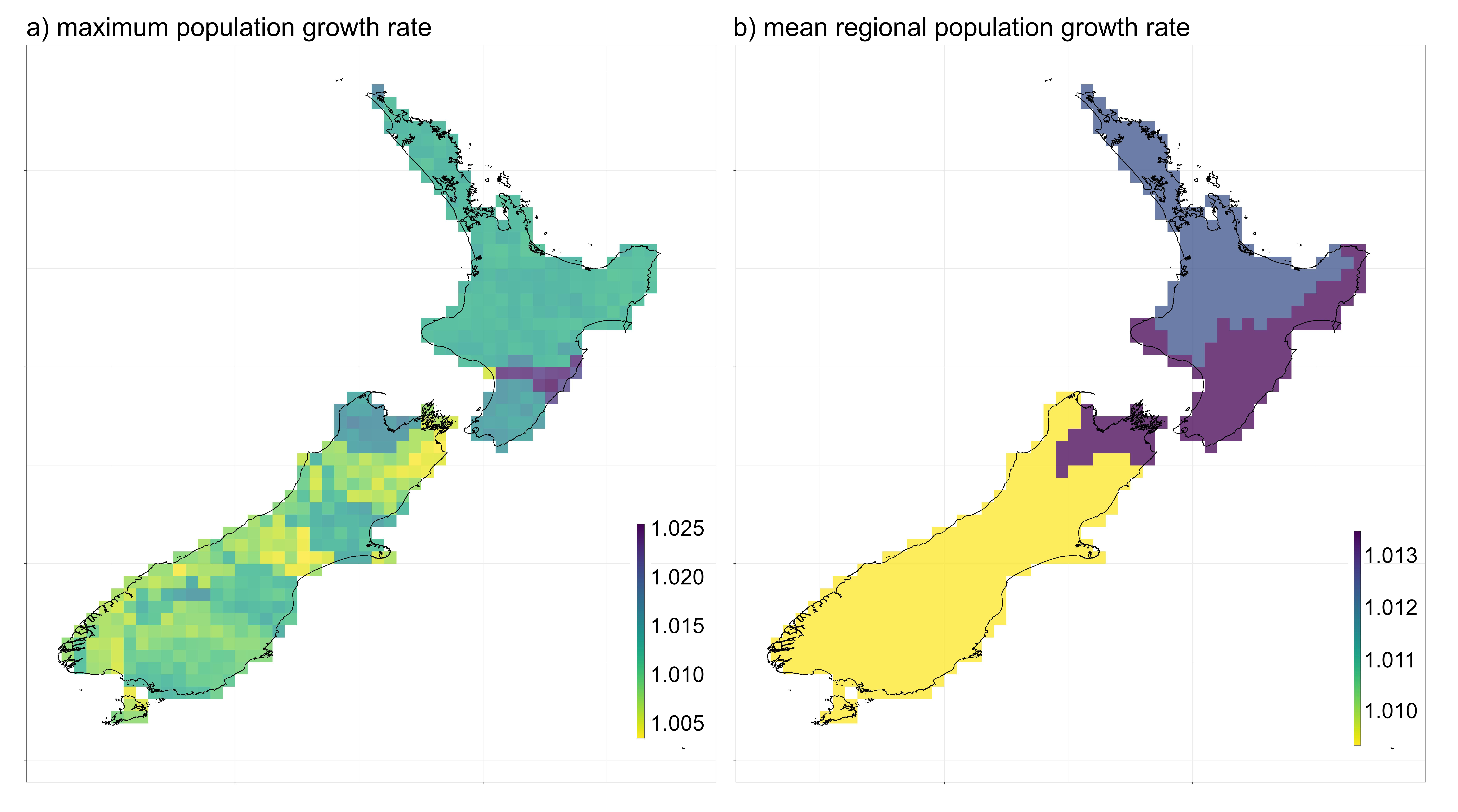

**Supplementary Figure 5:** a) Maximum annual population growth rates across New Zealand generated by the interaction between the population growth function and the spatial template of relative population density. When averaged according to economic subregions broadly consistent with those defined by Brown and Crema (2019) (b) population growth on the North Island was consistently higher than that in the South Island.

Population Growth Rates

When appropriately integrated with the SEPM, growth rates at the population level (i.e. at each neighbourhood) were spatially heterogenous (Supplementary Figure 5). This spatial pattern emerged as a result of the human dispersal model preferentially growing different populations in proportion to the relative population template. When summarised by bioregions, the population growth rates generated by our model were similar to those identified by Brown and Crema (2019), with highest rates of growth on the North Island, however our model suggests that maximal growth rates were highest in areas analogous to the Central economic subregion defined by Brown and Crema (2019) (approximately 1.013), followed by the Central economic subregion (approximately 1.012). The Southern economic subregion had much lower growth rates (approximately 1.01; Supplementary Figure 5)

Pattern oriented modelling

Bayes Factors indicated that the parameters were generally stable across all the iterations of the model, with the exception of the foraging radius of the Māori populations (Supplementary Table 2). This altered substantially between the iteration of the model using uninformed priors and the simulations selected by ABC. Following the first resampled iteration of the model, using informed parameter distributions, all parameters remained stable for all simulations. We found no significant differences in the Euclidean distances from the empirical targets between the first and second resampled iterations (t_25003_ = -49.5; p = 1). The credible parameter estimates that resulted from this process are summarised in Table 1.

**Supplementary Table 2:** Bayes Factors of resulting from tests of reiterations of the Māori expansion models to refine parameter space. The 1^st^ and 2^nd^ iterations result from informed priors drawn from the most accurate simulations identified by Approximate Bayesian Computation. Resampling was finalised at the 2^nd^ iteration following the return of a stable posterior predictive check.

| **Parameter** | **Uninformed priors** | **1^st^ iteration** | **2^nd^ iteration** |
| --- | --- | --- | --- |
| human_founding_population | 0 | 0 | 0 |
| human_growth_rate | 0 | 0 | 0 |
| human_colonisation_time | 0 | 0 | 0 |
| human_foraging_distance | 0 | 0 | 0 |
| human_min_neighbourhood | 53.8 | 0 | 0 |

SEPM Sensitivity Analyses

The simulations were sensitive to both the growth function (Supplementary Figure 6) and the number of founding events that constituted the founding population (Supplementary Figure 7). Neither of the alternative modelled scenarios could meet the empirical targets as well as the optimised simulations that we have reported here, but they could potentially be optimised to do so. Regardless, however, the spatial pattern of arrival did not differ substantially between the different population growth models (RMSE = 17.9; Supplementary Figure 6), with arrival at each grid cell following a consistent pattern and remaining within the error band established in the literature (G. L. Perry et al., 2014; Wilmshurst et al., 2008). The number of colonisation events did substantially alter the timing of colonisation (Supplementary Figure 7), where more numerous, smaller colonisation events resulted in much slower population growth and dispersal across the archipelago, and greater divergence between the models (RMSE (2 events) = 31.4; RMSE (5 events) = 110.0; RMSE (10 events) = 163.1). Although these structural changes have not been optimised through reiterations of ABC, it seems unlikely that multiple colonisation events modelled in this way would result in as accurate a match of the archaeological evidence as a small or even singular colonisation event, consistent with other reports (Walter et al., 2017).

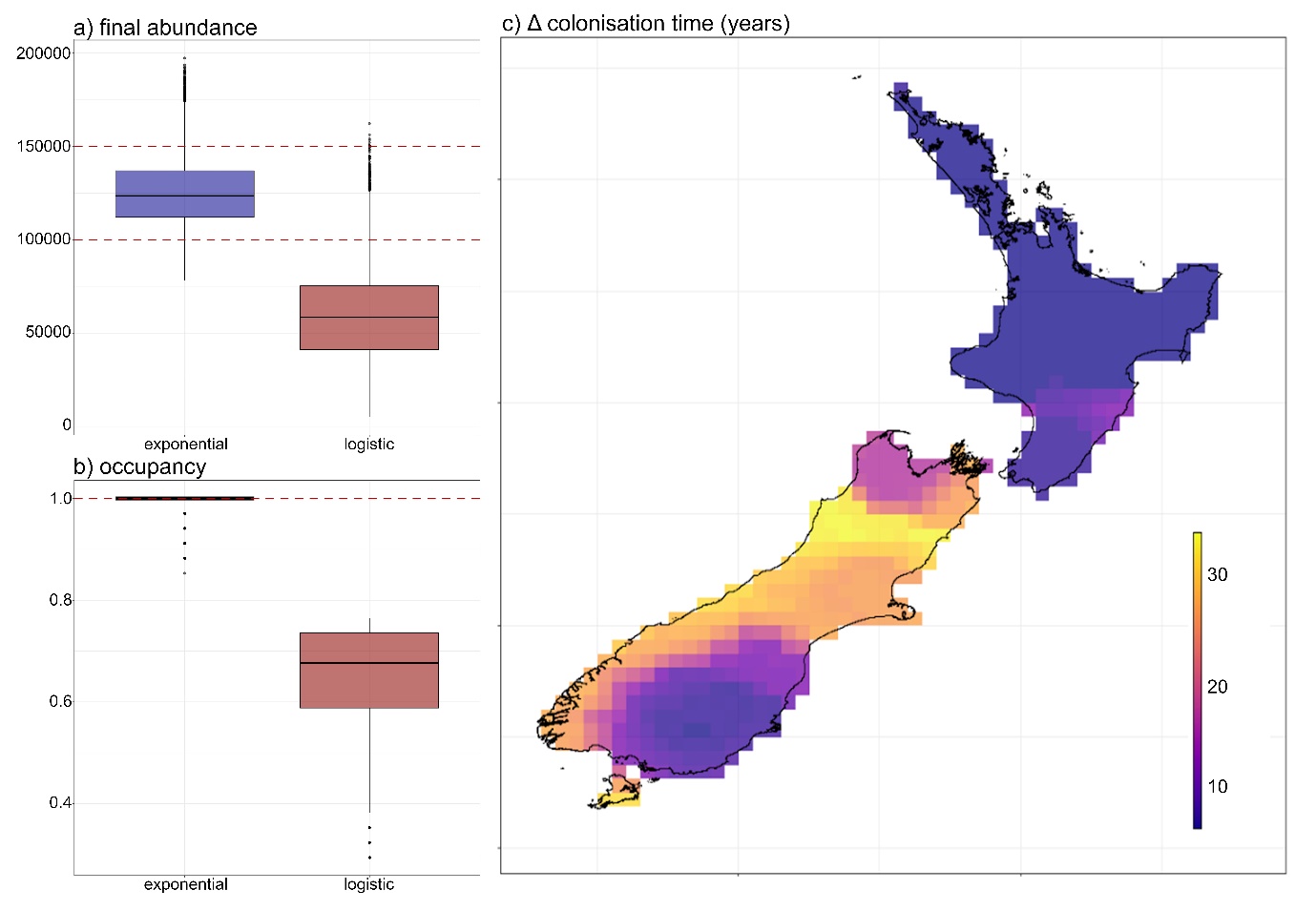
**Supplementary Figure 6:** Difference in model performance of simulations characterised by an exponential population growth rate (blue) and a logistic population growth rate (red) against **a)** a target of final abundance and **b)** a target of evidence of occupancy. Dashed red lines indicate the target range. **c)** The difference in mean weighted colonisation timing across New Zealand compared with the exponential growth model indicates errors that fall within the colonisation window.

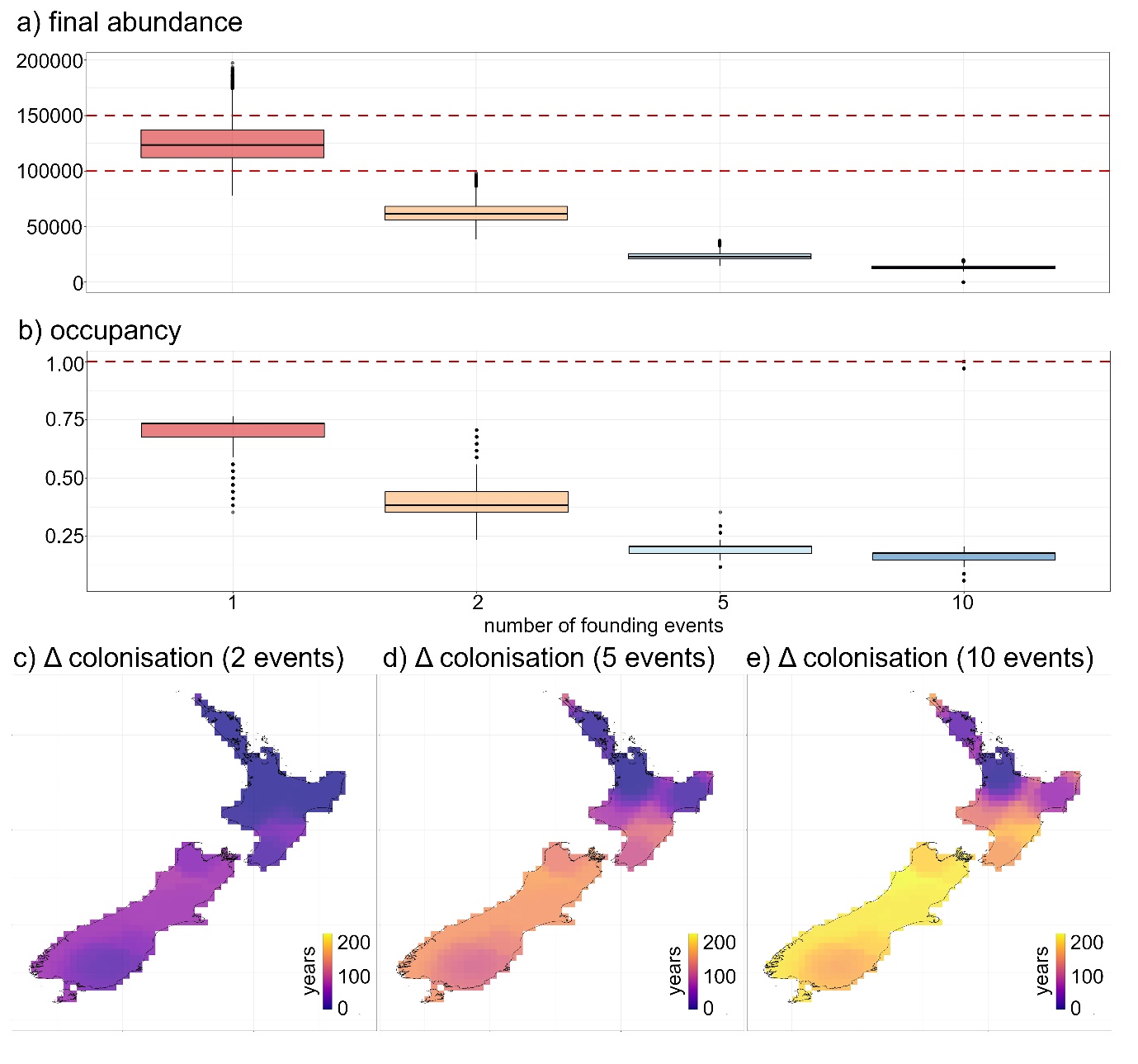

**Supplementary Figure 7:** Difference in model performance of simulations characterised by an increasing number of colonisation events against **a)** a target of final abundance and **b)** a target of evidence of occupancy. Dashed red lines indicate the target range. The difference in mean weighted colonisation timing across New Zealand comparing **c)** 2 founding events, **d)** 5 founding events and **e)** indicating increasing error in time of colonisation across New Zealand with an increasing number of colonisation events.

**REFERENCES**

Adams, B., White, A., & Lenton, T. M. (2004). 2004. An analysis of some diverse approaches to modelling terrestrial net primary productivity. *Ecological Modelling, 177*, 353-391.

Akçakaya, H. R. (1998). *RAMAS GIS: Linking Landscape Data with Population Viability Analysis (version 3.0).* Setauket, New York State, USA: Applied Biomathematics.

Allen, M. W. (2016). Food, fighting, and fortifications in pre-European New Zealand: beyond the ecological model of Maori warfare. In A. M. VanDerwarker & G. D. Wilson (Eds.), *The Archaeology of Food and Warfare - Food Insecurity in Prehistory* (pp. 41-59). Cham: Springer.

Baty, F., Ritz, C., Charles, S., Brutsche, M., Flandrois, J.-P., & Delignette-Muller, M.-L. (2015). A toolbox for nonlinear regression in R: The package nlstools. *Journal of Statistical Software, 66*, 1-21.

Bentley, G. R. (1985). Hunter-gatherer energetics and fertility: a reassessment of the !Kung San. *Human Ecology, 13*, 79-109.

Binford, L. R. (2001). *Constructing Frames of Reference: An Analytical Method for Archaeological Theory Building Using Ethnographic and Environmental Data Sets.* Berkeley, USA: University of California Press.

Bjornstad, O. N., & Falck, W. (2001). Nonparametric spatial covariance functions: estimation and testing. *Environmental and Ecological Statistics, 8*, 53-70.

Booker, D. J. (2015). *Hydrological indices for national environmental reporting. NIWA Client Report CHC2015-015.* Christchurch, New Zealand: National Institute of Water & Atmospheric Research.

Boyce, M. S., Vernier, P. R., S.E., N., & Schmiegelow, F. K. A. (2002). Evaluating resource selection functions. *Ecological Modelling, 157*, 281-300.

Brown, A. A., & Crema, E. R. (2019). Māori population growth in pre-contact New Zealand: Regional population dynamics inferred from summed probability distributions of radiocarbon dates. *The Journal of Island and Coastal Archaeology, 14*, 1-19.

Burnham, K. P., & Anderson, D. R. (2002). *Model Selection and Multimodel Inference: a Practical Information-theoretic Approach, 2^nd^ Edition.* New York: Springer.

Crespi, C. M., & Boscardin, W. J. (2009). Bayesian model checking for multivariate outcome data. *Computational Statistics & Data Analysis, 53*, 3765-3772.

Csilléry, K., Blum, M. G. B., Gaggiotti, O. E., & François, O. (2010). Approximate Bayesian Computation (ABC) in practice. *Trends in Ecology and Evolution, 25*, 410-418.

Csilléry, K., François, O., & Blum, M. G. B. (2012). abc: an R package for approximate Bayesian computation (ABC). *Methods in Ecology and Evolution, 3*, 475-479.

Di Cola, V., Broennimann, O., Petitpierre, B., Breiner, F. T., D'amen, M., Randin, C., . . . Pellissier, L. (2017). ecospat: an R package to support spatial analyses and modeling of species niches and distributions. *Ecography, 40*, 774-787.

Droogers, P., & Allen, R. G. (2002). Estimating reference evapotranspiration under inaccurate data conditions. *Irrigation and Drainage Systems, 16*, 33-45.

Elith, J., Leathwick, J. R., & Hastie, T. (2008). A working guide to boosted regression trees. *Journal of Animal Ecology, 77*, 802-813.

Fordham, D. A., Akçakaya, H. R., Brook, B. W., Rodríguez, A., Alves, P. C., Civantos, E., . . . Araujo, M. B. (2013). Adapted conservation measures are required to save the Iberian lynx in a changing climate. *Nature Climate Change, 3*, 899-903.

Fordham, D. A., Bertelsmeier, C., Brook, B. W., Early, R., Neto, D., Brown, S. C., . . . Araújo, M. B. (2018). How complex should models be? Comparing correlative and mechanistic range dynamics models. *Global Change Biology, 24*, 1357-1370.

Fordham, D. A., Brown, S. C., Akçakaya, H. R., Brook, B. W., Haythorne, S., Manica, A., . . . Nogués‐Bravo, D. (2021). Humans hastened the range collapse and extinction of woolly mammoth. . *Ecology Letters, Accepted*.

Fordham, D. A., Haythorne, S., & Brook, B. W. (2016). Sensitivity Analysis of Range Dynamics Models (SARDM): Quantifying the influence of parameter uncertainty on forecasts of extinction risk from global change. *Environmental Modelling & Software, 83*, 193-197.

Fordham, D. A., Haythorne, S., Brown, S. C., Buettel, J. C., & Brook, B. W. (2021). poems: R package for simulating species’ range dynamics using pattern-oriented validation. *Methods in Ecology and Evolution, 12*, 2364-2371.

Fordham, D. A., Resit Akçakaya, H., Araújo, M. B., Elith, J., Keith, D. A., Pearson, R., . . . Tozer, M. (2012). Plant extinction risk under climate change: are forecast range shifts alone a good indicator of species vulnerability to global warming? *Global Change Biology, 18*, 1357-1371.

Fordham, D. A., Saltré, F., Haythorne, S., Wigley, T. M., Otto‐Bliesner, B. L., Chan, K. C., & Brook, B. W. (2017). PaleoView: a tool for generating continuous climate projections spanning the last 21 000 years at regional and global scales. *Ecography, 40*, 1348-1358.

Fordham, D. A., Wigley, T. M., & Brook, B. W. (2011). Multi‐model climate projections for biodiversity risk assessments. *Ecological Applications, 21*, 3317-3331.

Furey, L. (2006). *Maori Gardening: An Archaeological Perspective.* Wellington: Department of Conservation.

Gelman, A., Hwang, J., & Vehtari, A. (2014). Understanding predictive information criteria for Bayesian models. *Statistics and Computing, 24*, 997-1016.

Giampoudakis, K., Marske, K. A., Borregaard, M. K., Ugan, A., Singarayer, J. S., Valdes, P. J., . . . Nogués‐Bravo, D. (2017). Niche dynamics of Palaeolithic modern humans during the settlement of the Palaearctic. *Global Ecology and Biogeography, 26*, 359-370.

Gigot, C. (2018). epiphy: Analysis of Plant Disease Epidemics. R package version 0.3.4. Vienna University of Economics and Business: The Comprehensive R Archive Network.

Goldberg, A., Mychajliw, A. M., & Hadly, E. A. (2016). Post-invasion demography of prehistoric humans in South America. *Nature, 532*, 232-235.

Greenwell, B., Boehmke, B., J., C., & Developers, G. (2019). gbm: Generalized Boosted Regression Models version 2.1.5. <https://CRAN.R-project.org/package=gbm>: The Comprehensive R Archive Network.

Gregory, S. D., Ancrenaz, M., Brook, B. W., Goossens, B., Alfred, R., Ambu, L. N., & Fordham, D. A. (2014). Forecasts of habitat suitability improve habitat corridor efficacy in rapidly changing environments. *Diversity and Distributions, 20*, 1044-1057.

Harris, I., Osborn, T. J., Jones, P., & Lister, D. (2020). Version 4 of the CRU TS monthly high-resolution gridded multivariate climate dataset. *Scientific Data, 7*, 1-18.

Haythorne, S., Fordham, D. A., Brown, S. C., Buettel, J. C., & Brook, B. W. (2021). poems: pattern-oriented ensemble modeling system. The Comprehensive R Archive Network.

Hirzel, A. H., & Le Lay, G. (2008). Habitat suitability modeling and niche theory. *Journal of Applied Ecology, 45*, 1372-1381.

Hirzel, A. H., Le Lay, G., Helfer, V., Randin, C., & Guisan, A. (2006). Evaluating the ability of habitat suitability models to predict species presences. *Ecological Modelling, 199*, 142-152.

Holdaway, R. N., Allentoft, M. E., Jacomb, C., Oskam, C. L., Beavan, N. R., & Bunce, M. (2014). An extremely low-density human population exterminated New Zealand moa. *Nature Communications, 5*, 5436.

Holdaway, R. N., & Jacomb, C. (2000). Rapid extinction of the moas (Aves: Dinornithiformes): model, test, and implications. *Science, 287*, 2250-2254.

Irwin, G., Johns, D., Flay, R. G., Munaro, F., Sung, Y., & Mackrell, T. (2017). A review of archaeological Māori canoes (Waka) reveals changes in sailing technology and maritime communications in Aotearoa/New Zealand, AD 1300–1800. *Journal of Pacific Archaeology, 8*, 31-43.

Krug, R. M. (2014). TSS: Basic and extended TSS Statistics (True Skill Statistics) version 0.3-0. <https://rdrr.io/github/rkrug/TSS/man/TSS-package.html>.

Lieth, H. (1973). Primary production: terrestrial ecosystems. *Human Ecology, 1*, 303-332.

Liu, C., Berry, P. M., Dawson, T. P., & Pearson, R. G. (2005). Selecting thresholds of occurrence in the prediction of species distributions. *Ecography, 28*, 385-393.

Makowski, D., Ben-Shachar, M. S., & Lüdecke, D. (2019). bayestestR: Describing effects and their uncertainty, existence and significance within the Bayesian framework. *Journal of Open Source Software, 4*, 1541.

Mellin, C., Russell, B. D., Connell, S. D., Brook, B. W., & Fordham, D. A. (2012). Geographic range determinants of two commercially important marine molluscs. *Diversity and Distributions, 18*, 133-146.

Norton, J. (2015). An introduction to sensitivity assessment of simulation models. *Environmental Modelling & Software, 69*, 166-174.

Pearce, C. E., & Pearce, F. M. (2010). The context of field archaeology: The Maori pa. In C. E. Pearce & F. M. Pearce (Eds.), *Oceanic Migration: Paths, Sequence, Timing and Range of Prehistoric Migration in the Pacific and Indian Oceans* (pp. 217-228). Dordrecht: Springer.

Perry, G. L., Wheeler, A. B., Wood, J. R., & Wilmshurst, J. M. (2014). A high-precision chronology for the rapid extinction of New Zealand moa (Aves, Dinornithiformes). *Quaternary Science Reviews, 105*, 126-135.

Perry, J. N. (1995). Spatial analysis by distance indices. *Journal of Animal Ecology, 64*, 303-314.

Perry, J. N. (1998). Measures of spatial pattern for counts. *Ecology*, 1008-1017.

Perry, J. N., & Hewitt, M. (1991). A new index of aggregation for animal counts. *Biometrics, 47*, 1505-1518.

Pool, D. I. (1991). *Te iwi Maori: a New Zealand Population, Past, Present and Projected.* Auckland, New Zealand: Auckland University Press.

Potts, J. M., & Elith, J. (2006). Comparing species abundance models. *Ecological Modelling, 199*, 153–163.

Prowse, T. A., Bradshaw, C. J., Delean, S., Cassey, P., Lacy, R. C., Wells, K., . . . Brook, B. W. (2016). An efficient protocol for the global sensitivity analysis of stochastic ecological models. *Ecosphere, 7*, p.e01238.

Ridout, M., Demétrio, C. G., & Hinde, J. (1998). Models for count data with many zeros. *Proceedings of the XIXth International Biometric Conference 19*, 179-192.

Saltelli, A., Tarantola, S., & Campolongo, F. (2000). Sensitivity analysis as an ingredient of modeling. *Statistical Science, 15*, 377-395.

Tallavaara, M., Eronen, J. T., & Luoto, M. (2018). Productivity, biodiversity, and pathogens influence the global hunter-gatherer population density. *Proceedings of the National Academy of Sciences, 115*, 1232-1237.

Timmermann, A., & Friedrich, T. (2016). Late Pleistocene climate drivers of early human migration. *Nature, 538*, 92-95.

Walker, R., Gurven, M., Hill, K., Migliano, A., Chagnon, N., De Souza, R., . . . Kramer, K. (2006). Growth rates and life histories in twenty‐two small‐scale societies. *American Journal of Human Biology, 18*, 295-311.

Walter, R., Buckley, H., Jacomb, C., & Matisoo-Smith, E. (2017). Mass migration and the Polynesian settlement of New Zealand. *Journal of World Prehistory, 30*, 351-376.

Wilmshurst, J. M., Anderson, A. J., Higham, T. F., & Worthy, T. H. (2008). Dating the late prehistoric dispersal of Polynesians to New Zealand using the commensal Pacific rat. *Proceedings of the National Academy of Sciences, 105*, 7676-7680.

Winder, L., Alexander, C., Griffiths, G., Holland, J., Woolley, C., & Perry, J. (2019). Twenty years and counting with SADIE: Spatial Analysis by Distance Indices software and review of its adoption and use. *Rethinking Ecology, 4*, 1-16.

Zealand, L. I. N. (2018). *NZ Coastlines (Topo, 1:50k)*.

Zealand, L. R. N. (2010). *LENZ - 25 metre Slope*.
